# A cortical signature of very preterm birth across development and its association with neurodevelopmental outcomes

**DOI:** 10.1101/2025.10.17.683086

**Authors:** Kaili Liang, Yourong Guo, Logan Z. J. Williams, Renato Besenczi, Zeyuan Sun, Marguerite Leoni, Qiang Ma, Daniel Rueckert, Paola Dazzan, A. David Edwards, Emma C. Robinson, Chiara Nosarti

## Abstract

Gestational age (GA) is a key determinant of neurodevelopment, and individuals born very preterm (VPT; ≤ 32 weeks’ gestation) are at increased risk of cognitive and behavioural difficulties across the lifespan. A better understanding of the impact of VPT birth on cortical maturation trajectories may provide mechanistic insight into the origins of these neurodevelopmental sequelae. Here we compared cortical morphology between VPT individuals and full-term controls in three datasets spanning the neonatal period, childhood and adulthood. We identified a consistent cortical signature of VPT birth, characterized by reduced surface area and cortical folding in the frontal, temporal, parietal and insular regions, which were observed across development. Reduced surface area and cortical folding in the neonatal period were associated with clinical factors and poorer motor outcomes in toddlerhood. Furthermore, longitudinal analyses in a VPT cohort showed that greater cortical expansion from the neonatal period to childhood was associated with higher childhood IQ. Together, these findings demonstrate persistent cortical morphological alterations following VPT birth and their relevance to long-term neurodevelopmental outcomes.

## 1 Introduction

From approximately 20 weeks of gestation until birth and into the early postnatal period, the cerebral cortex undergoes rapid exponential expansion, characterized by marked increases in cortical folding and surface area (Bethlehem et al., 2022; Kostovic et al., 2019). These processes are critical for supporting motor, cognitive and behavioural development in infancy and childhood (Houwen et al., 2016; Shaw et al., 2006). Very preterm birth (VPT; ≤ 32 weeks’ gestation), occurring during this critical neurodevelopmental window, is therefore widely understood to disrupt typical cortical maturation, resulting in long-lasting alterations in cortical organization (de Vareilles et al., 2023; Garcia et al., 2018; Kelly et al., 2023; Kelly et al., 2024). Moreover, compared with full-term individuals, those born VPT are at increased risk of suboptimal neurodevelopmental outcomes, including early-life motor and cognitive impairments and increased neuropsychiatric symptoms throughout childhood and beyond (Bröring et al., 2018; Brydges et al., 2018; Lean et al., 2018; Nosarti et al., 2012). As a result, VPT birth presents a valuable model for investigating how early changes in cortical development impact subsequent cognitive and behavioural outcomes, which can provide insights into the neurobiological mechanisms underlying neurodevelopmental disorders. Advances in neuroimaging, particularly structural MRI, have enabled the characterisation of cortical features that may provide mechanistic insights into these developmental pathways.

Structural MRI has been widely used to quantify cortical morphology, including surface area, reflecting the number of radial cortical columns; sulcal depth, a measure of cortical folding complexity; and cortical thickness, commonly interpreted as a proxy for cytoarchitectural properties such as synaptic density and dendritic arborization (Chiarello et al., 2016). Numerous studies have reported significant cortical morphological alterations in individuals born preterm (< 37 weeks’ gestation), including those born VPT, compared with full-term individuals (Bonthrone et al., 2025; Dimitrova et al., 2021; Engelhardt et al., 2015; Kelly et al., 2023; Kline et al., 2019; Papini et al., 2020; Thalhammer et al., 2024; Zhang et al., 2015). Most evidence suggests reduced cortical surface area in temporal, frontal, and parietal regions in preterm or VPT individuals (Engelhardt et al., 2015; Kelly et al., 2023; Kline et al., 2019; Thalhammer et al., 2024; Zhang et al., 2015). However, a recent longitudinal study reported accelerated surface area expansion in frontal, temporal, and supramarginal/inferior parietal regions from term-equivalent age to 9–10 years in VPT children, resulting in surface areas comparable to those of full-term children, and suggesting a potential "catch-up" effect (Gorham et al., 2024). In contrast, findings regarding cortical thickness are more heterogeneous. At term-equivalent age, studies have reported overall increased cortical thickness in preterm neonates relative to full-term controls (Dimitrova et al., 2021; Thalhammer et al., 2024). However, by childhood or adulthood, prior work has documented both regional increases and decreases in cortical thickness (Kelly et al., 2023; Nam et al., 2015; Sripada et al., 2018), as well as predominantly thinner cortices in frontal and temporal regions (Thalhammer et al., 2024). Taken together, these mixed findings indicate that it remains unclear whether a distinct cortical footprint associated with VPT birth persists across development.

Most studies of brain-behaviour relationships have focused on cross-sectional associations between cortical morphological alterations and cognitive outcomes in preterm children and adults (Inder et al., 2023; Papini et al., 2020; Sripada et al., 2018; Thalhammer et al., 2024), suggesting that reduced cortical surface area and folding are linked to cognitive impairments (Papini et al., 2020; Thalhammer et al., 2024). However, infancy represents a critical period for the development of motor, language and cognitive functions that support higher-order cognitive, behavioral and emotional regulation later in life (Houwen et al., 2016). Accordingly, longitudinal investigations of neonatal cortical morphology and subsequent neurodevelopment are essential to elucidate the neurobiological mechanisms linking early cortical alterations to later neurodevelopmental impairments. Importantly, examining developmental trajectories may provide additional insights beyond early structural differences alone. For example, a longitudinal study of VPT individuals demonstrated that atypical cortical development from infancy to childhood was associated with poorer language and memory outcomes in childhood (Kelly et al., 2024), highlighting the importance of understanding how patterns of cortical maturation may contribute to later cognitive functioning.

In this study, we firstly examined whether a distinct VPT cortical signature persisted across development by comparing cortical thickness, surface area and sulcal depth between VPT individuals and full-term controls in three cross-sectional datasets spanning neonates, children and adults. In the neonatal dataset, we further investigated associations between neonatal cortical morphology, clinical risk factors and cognitive, language and motor outcomes at 18–24 months. These association analyses were repeated in an independent VPT infant cohort. Finally, in a longitudinal VPT cohort with imaging acquired at both the neonatal and childhood time points, we examined whether cortical expansion from infancy to childhood was associated with GA and childhood IQ.

## 2 Methods

### 2.1 Participants

The study design is shown in Figure 1. Three cross-sectional datasets were used to examine cortical alterations in VPT individuals: 541 full-term (FT) and 81 VPT neonates from the Developing Human Connectome Project (dHCP) (Edwards et al., 2022), 81 FT and 158 VPT children from the Brain, Immunity, and Psychopathology following Very Preterm Birth (BIPP) study (Sun et al., 2025), and 95 FT and 118 VPT adults from the University College London Hospital (UCLH) cohort (Papini et al., 2020). Across these datasets, FT participants’ gestational age (GA) ranged between 37 and 42 weeks, and VPT participants were born at <33 weeks’ gestation. To investigate associations between neonatal cortical morphology, clinical risk factors and later neurodevelopment, we further analysed 697 preterm and full-term neonates (GA range: 23–42 weeks) from the dHCP; and analyses were replicated in an independent sample of 414 VPT neonates (<33 weeks’ GA) from the Evaluation of Preterm Imaging (ePrime) study (Edwards et al., 2018).

**Figure 1.**
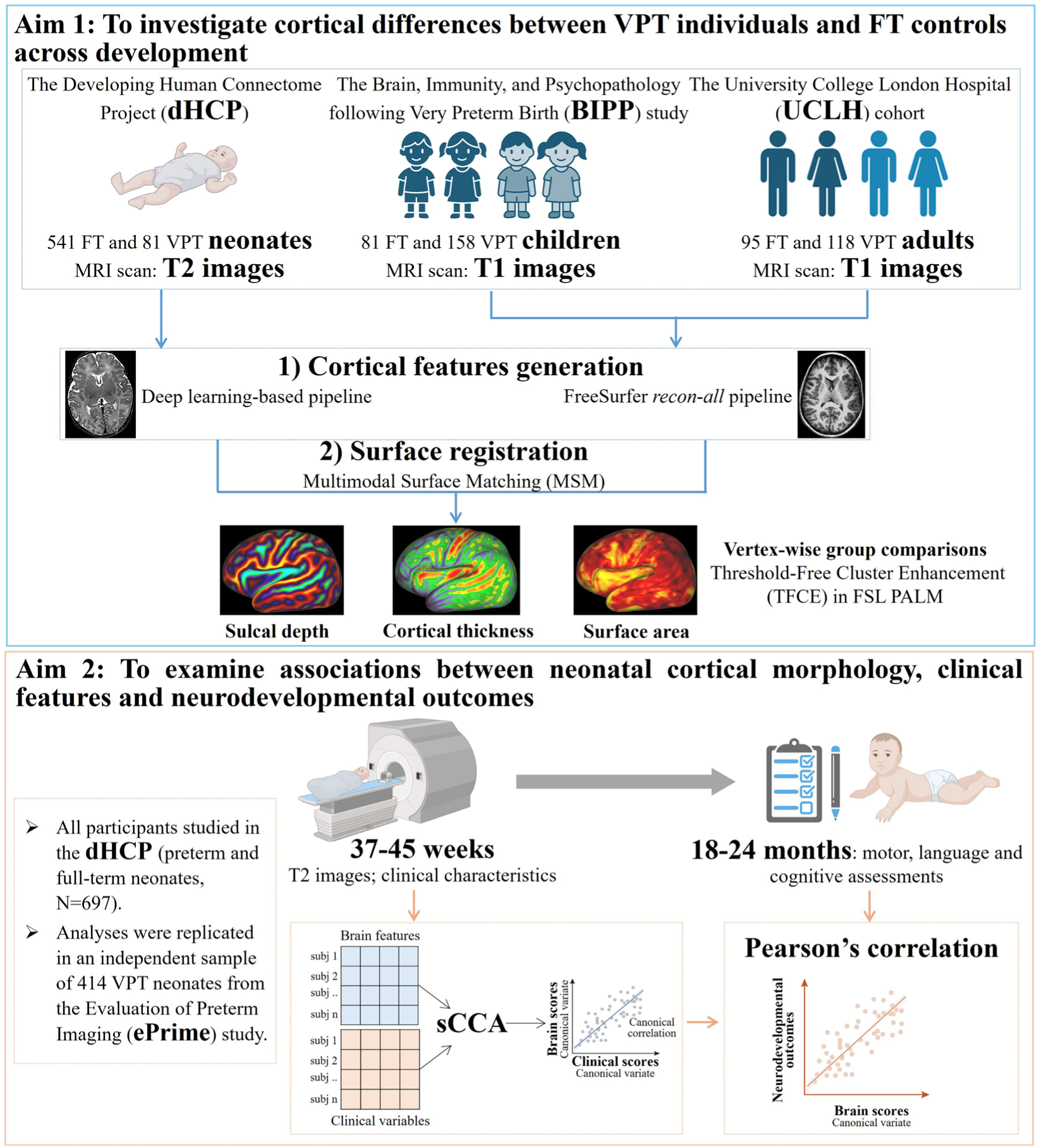
Study design. VPT: very preterm; FT: full-term; sCCA: sparse canonical correlation analysis.

The BIPP study represents the childhood follow-up of the ePrime study (Edwards et al., 2018). Among the 414 VPT participants with neonatal cortical surface data, 148 also underwent MRI in childhood and were included in the longitudinal analyses of cortical expansion in relation to GA and childhood IQ.

All studies received ethical approval from the relevant Research Ethics Committee. Details of ethics approvals, recruitment periods and sites, inclusion and exclusion criteria for each dataset are provided in Supplementary Materials, *Participants* section.

### 2.2 Socio-demographic, clinical and behavioural measures

Socio-demographic variables collected in all datasets included gestational age at birth, age at scan, sex, and socio-economic status (SES). SES was assessed using the Index of Multiple Deprivation (IMD) in the dHCP, BIPP, and ePrime cohorts, an area-level composite measure based on residential postcodes incorporating income, employment, education, crime, and health. The 2015 IMD (https://www.gov.uk/government/statistics/english-indices-of-deprivation-2015) was used for the dHCP and BIPP datasets, while the 2010 IMD (https://tools.npeu.ox.ac.uk/imd/) was applied to the ePrime due to earlier data collection. In the UCLH cohort, SES was determined using the UK Stationery Office Occupational Classification and dichotomized into high (managerial/professional) and low (all other) categories (Censuses et al., 1995).

Participants’ available clinical characteristics are reported in Supplementary Materials, *Clinical Features* section. Clinical factors explored in relation to neonatal cortical morphology in dHCP and ePrime included gestational age and birth weight.

Neurodevelopmental outcomes at 18–24 months were assessed in dHCP and ePrime using the Bayley Scales of Infant and Toddler Development–Third Edition (BSID-III) (Wechsler, 2003). Age-standardized cognitive, language, and motor subscale scores were used in our analyses. Childhood IQ in BIPP was assessed using the Wechsler Intelligence Scale for Children (WISC-IV) (Petermann & Petermann, 2011), and age-normed full-scale IQ scores were used in the analyses.

### 2.3 MRI acquisition and processing

For the dHCP and ePrime cohorts, all participants underwent MRI at term-equivalent age (37–45 weeks postmenstrual age). In BIPP, MRI was acquired during childhood (7–13 years), and in the UCLH cohort in adulthood (23–45 years). Cortical morphological analyses were performed using T1-weighted images in the BIPP and UCLH datasets. For neonates in the dHCP and ePrime datasets, analyses were conducted on T2-weighted images due to the reversal of grey-white matter contrast in neonatal MRI relative to that observed in children and adults (Edwards et al., 2022). Detailed MRI acquisition parameters are provided in the *MRI acquisition* section of the Supplementary Materials.

Surface-based analysis using structural MRI has been widely advocated for studying the cerebral cortex due to the fact that it regularises image processing with respect to geodesic distances along the cortical sheet, which are more neurobiologically meaningful than the three-dimensional Euclidean distances used in volume-based analyses (Glasser et al., 2016; Glasser et al., 2013). We therefore utilized surface-based analyses, including surface reconstruction and registration, to investigate cortical morphology in this study.

#### 2.3.1 Surface extraction and feature generation

For neonates in dHCP and ePrime, surface reconstruction was performed using a novel deep learning-based (DL) pipeline (Ma et al., 2025). Briefly, the preprocessing of T2w images involved deep learning-based brain extraction and N4 bias field correction. Subsequently, an end-to-end learning-based approach for cortical surface reconstruction was applied to generate white matter and pial surfaces. Inflated surfaces were then derived from the white matter surface, with sulcal depth or estimates of mean convexity/concavity for each vertex calculated during the inflation process. Finally, spherical projection was achieved through learning-based spherical projection; specifically the mapping from white matter surface to the sphere was optimised using a spherical U-Net (Zhao et al., 2022), optimising the same losses used by the classical FreeSurfer approach: edge and area distortions (Fischl et al., 1999). Compared with the spherical multidimensional scaling (MDS) approach used in the previous dHCP pipeline (Makropoulos et al., 2018), this approach produces spherical representations with reduced edge and area distortions (Ma et al., 2025).

For children in BIPP and adults in UCLH cohort, the *recon-all* pipeline in FreeSurfer (version 6.0.0) was employed to produce surface meshes and cortical features (Fischl, 2012); this performs motion correction, brain extraction, intensity normalization, tissue segmentation, topology correction, surface extraction, inflation and spherical mapping. The ciftify_recon_all (https://edickie.github.io/ciftify/#/03a_cifti-for-your_recon_all) was then utilized to convert recon-all outputs into an HCP-style folder structure. This conversion was conducted to transform surface meshes and metrics into the GIFTI file format for visualization and further analyses (Dickie et al., 2019).

The cortical features used in this study include cortical thickness (calculated by averaging the bidirectional distances between the white matter and pial surfaces), sulcal depth (measured during the inflation process), and surface area (computed from the white matter surface). To correct for folding bias, cortical thickness and surface area were adjusted by regressing out cortical curvature, following the methodology of the HCP minimal preprocessing pipeline (Glasser et al., 2013). Quality control was performed on white matter and pial surfaces by visual inspection and were deemed usable.

#### 2.3.2 Surface registration in cross-sectional datasets

The purpose of surface registration is to align corresponding brain regions across individuals, enabling group-level comparisons at anatomically consistent locations within a common template space. In this study, registration was performed using Multimodal Surface Matching (MSM), a flexible spherical registration approach that enables accurate alignment of surfaces based on various features (Besenczi et al., 2024; Robinson et al., 2018; Robinson et al., 2014).

Here we employed MSMSulc, an MSM variant optimized for correspondence of sulcal depth, to register individual cortical surfaces to age-matched template spaces.

Template selection is a critical step before surface registration. Two key considerations are: (1) whether the template reflects the study population’s demographics and (2) the impact of spherical projection on data representation. For children and adults, one population-average template derived from young adult data, such as the MSMSulc HCP template, is commonly used, as human brain shape changes minimally after age 2 (Kostovic et al., 2019). In contrast, neonatal brains undergo rapid morphological changes in size and shape, requiring spatio-temporally evolving templates specific to each gestational week (Bozek et al., 2018; Williams et al., 2023). Since the spherical projection step in the dHCP deep learning pipeline is not compatible with that used in the previous dHCP pipeline, we first regenerated week-specific neonatal surface templates from the new surfaces using the template construction framework described by Bozek et al. (2018) and Williams et al. (2023) (Bozek et al., 2018; Williams et al., 2023). Detailed procedures are provided in the *Neonatal Template Generation* section of the Supplementary Materials. The newly generated week-specific dHCP templates with left–right vertex correspondence are shown in Supplementary Figures 1 and 2.

Neonatal cortical surfaces were aligned to their closest week-specific dHCP template using MSMSulc and subsequently propagated to a common week-40 reference space through template-to-template alignment. Children and adult cortical surfaces were aligned to the HCP template using MSMSulc. For cross-stage analyses, the dHCP week-40 template was registered to the HCP template following Williams et al. (2023) (Williams et al., 2023), enabling all neonatal surfaces to be aligned to the HCP template space. Further details of the registration procedure are provided in the *Surface Registration* section of the Supplementary Materials. Surface meshes and metrics were resampled using barycentric and adaptive barycentric interpolation, respectively. Surface registration quality was assessed by visual inspection using the Connectome Workbench (*wb_view*), by comparing template and template-registered sulcal depth maps with major anatomical landmarks. A detailed quality control protocol is provided in the publicly available MSM video tutorials (https://www.youtube.com/watch?v=rQQx0k1Sgdc).

#### 2.3.3 Longitudinal surface registration

To quantify longitudinal cortical growth from the neonatal period to childhood, we used anatomically constrained Multimodal Surface Matching (aMSM), an extension of the MSM framework that regularises deformations on the anatomical white matter surface rather than the spherical surface, providing biologically plausible estimates of longitudinal cortical expansion (Besenczi et al., 2024; Garcia et al., 2018; Robinson et al., 2018). This framework has been validated for both fetal-to-neonatal and neonatal-to-childhood longitudinal surface registration (da Silva et al., 2025; Garcia et al., 2018; Gorham et al., 2024). Longitudinal surface registration was performed using the same configuration and parameters as described by da Silva et al. (2025) (da Silva et al., 2025). Registration accuracy was visually assessed for each participant. Following longitudinal registration, the vertex-wise relative cortical expansion map was calculated for each participant as:

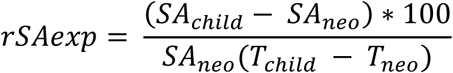

where *SA_neo_* and *SA_child_* denote the vertex-wise cortical surface area at the neonatal and childhood time points, respectively, and *T_neo_* and *T_child_* represent the corresponding scan ages (days). Therefore, *rSAexp* represents the percentage change in cortical surface area per day between the two scans.

### 2.4 Statistical analysis

#### 2.4.1 Group comparisons at each developmental stage

Cortical features, including cortical thickness, surface area and sulcal depth, were compared between VPT and FT groups at each developmental stage using surface-based, vertex-wise permutation testing with Threshold-Free Cluster Enhancement (TFCE) in FSL PALM (Smith & Nichols, 2009; Winkler et al., 2014). Analyses were conducted with age at scan, sex, total brain volume, and socioeconomic status as covariates. Family-Wise Error (FWE) corrections were applied to *-log(P)* values across three cortical features and two design contrasts (VPT > FT and VPT < FT). Statistical significance was defined as *-log(P)_FWE* > 1.6, corresponding to an adjusted *P* < 0.025 per hemisphere to account for bilateral comparisons.

#### 2.4.2 Group comparisons across developmental stages

The ComBat method (Fortin et al., 2018) was firstly employed to remove site-specific effects while preserving biological information, including sex, prematurity (VPT or FT), age at scan and total brain volume, at the level of vertex. This method has been widely applied in neuroimaging studies for data harmonization (Bedford et al., 2025; Kang et al., 2024; Keller et al., 2023). To standardize age at scan across different developmental stages, age in years was converted to weeks by multiplying by 52.18, which represents the average number of weeks in a calendar year (365.25/7). After harmonization, vertex-wise group comparisons between VPT and FT participants across the three datasets were performed using FSL PALM (Smith & Nichols, 2009; Winkler et al., 2014), controlling for sex, age at scan and total brain volume. Here, socioeconomic status was not included as a covariate due to differences in its assessment across datasets. Vertex-wise effect sizes (*Cohen’s d*) were calculated for significant regions of group comparison.

Next, regions exhibiting significant group differences were used as masks to extract individual-level average cortical feature values. We then compared these regional mean values between groups at each developmental stage, using scanning age, sex, total brain volume, and socioeconomic status as covariates.

#### 2.4.3 Associations between clinical factors, neurodevelopmental outcomes and cortical alterations

We employed sparse canonical correlation analysis (sCCA) (Avants et al., 2014; Smith et al., 2015; Witten et al., 2009) to examine associations between neonatal cortical morphology and clinical risk factors, following the method described by Xia et al,. (Xia et al., 2018). The effects of age at scan, sex, IMD, and brain volume were first regressed out from vertex-wise cortical feature values from significant regions identified across development. Residuals of cortical feature values, along with clinical factors (gestational age and birth weight), were used as inputs for sCCA. Elastic-net regularization, combining LASSO and ridge penalties, was applied to obtain sparsity and mitigate overfitting. A grid search was conducted to optimize regularization parameters by maximizing the correlation of the first canonical variate across 10 randomly resampled subsets, each comprising two-thirds of the dataset. Here the optimal regularisation parameters were estimated in dHCP and subsequently applied to the ePrime cohort without further tuning. Statistical significance was assessed via permutation testing (*n*=1000). To evaluate the stability of sCCA results, we performed 1000 bootstrapped resamples with correlations deemed stable if their 95% confidence intervals did not include zero.

The sCCA identifies canonical correlation modes, each comprising a pair of canonical variates. In this study, canonical variates derived from cortical variables are referred to as brain scores, and those derived from clinical factors as clinical scores. Brain and clinical loadings were quantified as correlation coefficients between individual variables and their corresponding canonical variates.

Pearson’s correlation was further used to examine relationships between brain scores from the above sCCA and age-normed BSID-III cognitive, language and motor scores, with scores adjusted for sex and socioeconomic status prior to analysis. Given that the dHCP included both preterm and full-term infants, group × brain score interactions with neurodevelopmental outcomes were tested prior to correlation analyses. Multiple comparisons were corrected using the Bonferroni method. Statistical analyses were conducted using R and Python.

To externally validate the associations between cortical alterations, clinical factors, and neurodevelopmental outcomes, brain and clinical loadings derived from the dHCP sCCA model were projected onto the ePrime dataset to generate corresponding brain and clinical scores. Associations between brain scores and neurodevelopmental outcomes were then evaluated. Reciprocal validation was performed by applying the loadings derived from the ePrime dataset to the dHCP cohort.

#### 2.4.4 Associations of cortical expansion from infancy to childhood with GA and childhood IQ within VPT individuals

Regions exhibiting significant differences in the group comparisons across development were used as masks to extract individual-level mean relative cortical expansion. Within VPT participants, associations of cortical expansion from infancy to childhood with GA and childhood full-scale IQ were assessed using Pearson’s correlations. Before analysis, cortical expansion values were adjusted for sex, relative brain volume growth from infancy to childhood, and socioeconomic status (IMD). Childhood full-scale IQ scores were adjusted for sex and IMD. Bonferroni correction was applied to control for multiple comparisons.

## 3 Results

Detailed demographic information for participants in the dHCP, ePrime, BIPP, and UCLH cohorts is reported in Table 1. Across the four datasets, one-way ANOVA within the VPT groups revealed an overall difference in gestational age (F = 2.87, P = 0.036); however, no pairwise comparisons remained significant after correction for multiple comparisons. No significant difference was observed in birth weight (F = 2.17, P = 0.090) within the VPT participants across the four datasets. In the two infant cohorts, 541/697 participants (77.6%) in dHCP (mean age 19.01 months) and 366/414 (88.4%) participants in ePrime (mean age 20.18 months) completed the follow-up BSID-III assessments. In ePrime, mean cognitive, language, and motor scores were 92.58, 90.92, and 94.99, respectively; corresponding scores in dHCP were 100.53, 97.97, and 101.61.

**Table 1.** Demographic characteristics of participants across the four study datasets.

|  | VPT group | FT group | P value |
| --- | --- | --- | --- |
| **Neonates: dHCP** | **N=81** | **N=541** | / |
| Gestational age at birth (weeks) | 29.07(2.77) | 39.90 (1.21) | <0.001 |
| Post-menstrual age at scan (weeks) | 41.48 (1.85) | 41.32 (1.72) | 0.475 |
| Sex (female) | 39 (48.15%) | 251 (46.40%) | 0.861 |
| Birth weight (kg) | 1.20 (0.43) | 3.38 (0.51) | <0.001 |
| IMD | 14792.22 (8555.26) | 13613.98 (7327.25) | 0.242 |
| **Neonates: ePrime** | **N=414** | / | / |
| Gestational age at birth (weeks) | 29.80 (2.26) | / | / |
| Post-menstrual age at scan (weeks) | 42.20 (1.62) | / | / |
| Sex (female) | 208 (50.24%) | / | / |
| Birth weight (kg) | 1.32 (0.39) | / | / |
| IMD | 20.13 (11.74) | / | / |
| **Children: BIPP** | **N=158** | **N=81** | / |
| Gestational age at birth (weeks) | 29.78 (2.29) | 40.17(1.21) | <0.001 |
| Age at scan (years) | 10.10 (1.51) | 9.07 (1.08) | <0.001 |
| Sex (female) | 81 (51.27%) | 44 (54.32%) | 0.756 |
| Birth weight (kg) | 1.32 (0.39) | / | / |
| IMD | 18506.02 (8885.50) | 19707.53 (9666.41) | 0.352 |
| **Adults: UCLH cohort** | **N=118** | **N=95** | / |
| Gestational age at birth (weeks) | 29.42 (2.13) | / | / |
| Age at scan (years) | 31.25 (2.35) | 30.06 (4.63) | 0.025 |
| Sex (female) | 52 (44.07%) | 52 (54.74%) | 0.158 |
| Birth weight (kg) | 1.32 (0.36) | / | / |
| SES (low) | 67 (56.78%) | 53 (55.79%) | 0.995 |

Data were shown as mean (standard deviation) or number (percentage). dHCP: the Developing Human Connectome Project; BIPP: the Brain, Immunity, and Psychopathology following Very Preterm Birth; UCLH: the University College London Hospital; VPT: very preterm; FT: full term; IMD: index of multiple deprivation; SES: socio-economic status; /: not available.

### 3.1 Cortical alterations in VPT participants at each developmental stage

Cortical differences between VPT and FT participants at each developmental stage are shown in Figure 2. VPT neonates exhibited shallower sulci in the bilateral insular, orbitofrontal, and intraparietal regions, as well as the right superior temporal and collateral sulci. Flatter gyri in VPT neonates were observed in the bilateral superior and middle temporal gyri, supramarginal gyri, precuneus, and the right inferior parietal lobule. For surface area, VPT neonates showed reductions in multiple regions, including the bilateral superior and inferior frontal regions; orbitofrontal and insular regions; superior and middle temporal regions; supramarginal region; banks of the superior temporal sulcus; superior and inferior parietal regions; fusiform and parahippocampal regions; and the left anterior cingulate cortex. In contrast, increased surface area was found in the bilateral medial occipital regions. Regarding cortical thickness, VPT neonates had thicker cortices in the bilateral insular regions, right central sulcus, right superior temporal region, and right medial occipital region, while thinner cortices were observed in the bilateral superior frontal, orbitofrontal, precentral, paracentral, postcentral, superior parietal, and precuneus regions.

**Figure 2.**
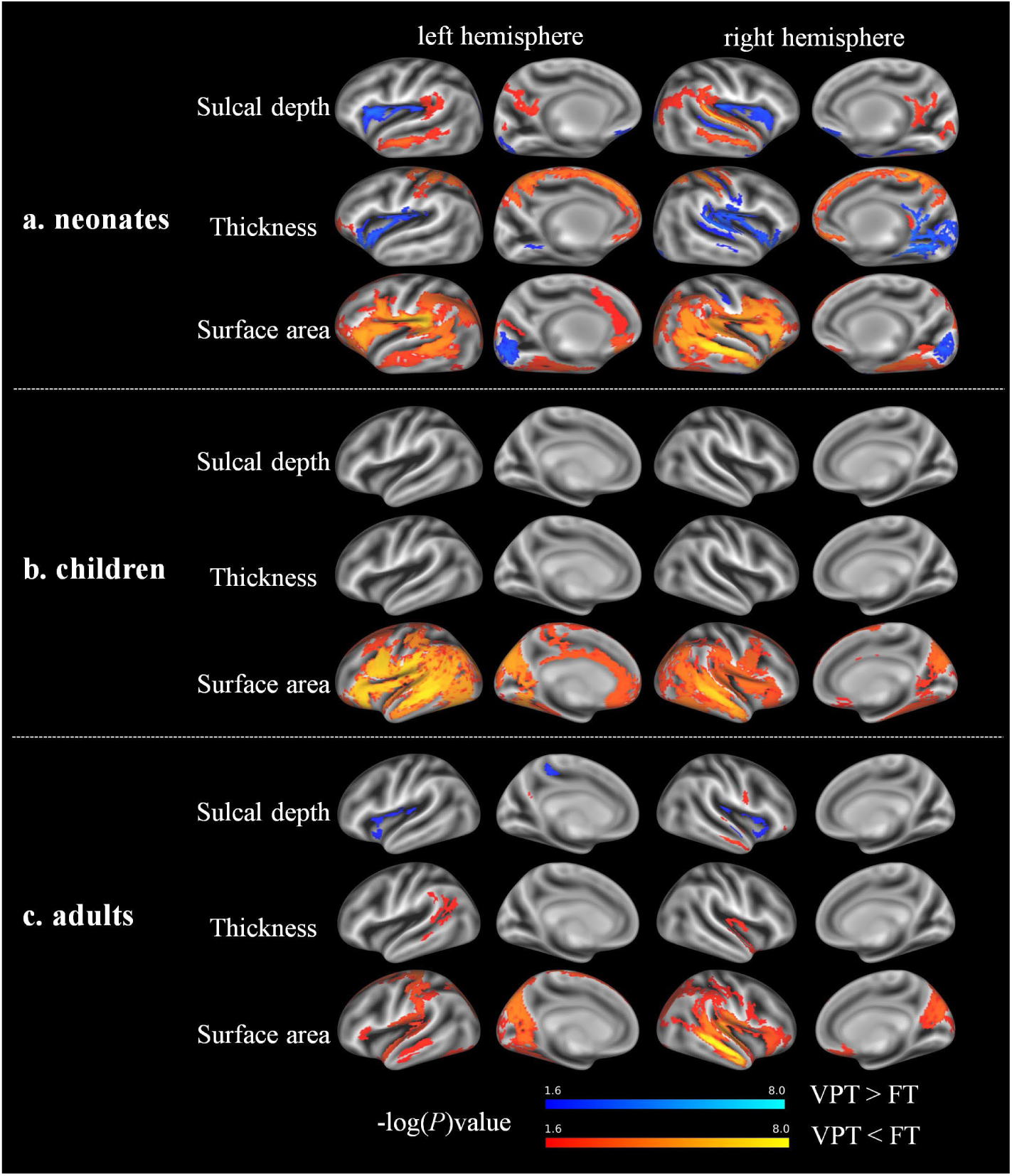
Group comparisons between very preterm (VPT) and full-term (FT) individuals at each developmental stage. Statistically significant regions are shown in blue–light blue (higher values in VPT than in FT) and red–yellow (reduced values in VPT compared with FT). Significance was determined using family-wise error (FWE) correction across three cortical features and two contrasts (VPT > FT, VPT < FT), with a threshold of –log(P) > 1.6 (adjusted P < 0.025 per hemisphere). Unlike surface area and cortical thickness maps, which contain only positive values, sulcal depth maps include negative values (representing sulci) and positive values (representing gyri); thus, higher values indicate shallower sulci and more prominent gyri, while lower values indicate deeper sulci and flatter gyri.

In childhood, VPT participants showed widespread surface area reductions in the bilateral superior, caudal middle, and inferior frontal regions; orbitofrontal and insular regions; superior, middle, and inferior temporal regions; supramarginal regions; banks of the superior temporal sulcus; superior and inferior parietal regions; lateral and medial occipital regions; posterior precuneus; fusiform; and parahippocampal regions. Additionally, reductions were observed in the left precentral, paracentral, postcentral and cingulate regions. No significant differences were found in sulcal depth or cortical thickness.

In adulthood, VPT participants exhibited shallower sulci in the bilateral insular regions and a flatter right middle temporal gyrus. Surface area reductions in VPT adults were observed in the bilateral inferior frontal, insular, precentral and postcentral regions; superior and middle temporal regions; supramarginal and superior parietal regions; posterior precuneus; medial occipital regions; as well as the left superior frontal and right orbitofrontal regions. In terms of cortical thickness, VPT adults demonstrated thinner cortices in the left supramarginal region, banks of the superior temporal sulcus, and the right superior temporal and insular region.

### 3.2 A consistent VPT cortical signature across development

The statistically significant results of group comparisons across development are shown in Figure 3a. Compared to full-term controls, VPT individuals exhibited shallower sulci in the bilateral insular, orbitofrontal, and intraparietal regions; superior temporal sulcus; posterior cingulate sulcus; collateral sulcus; as well as the right inferior frontal sulcus. Flatter gyri in VPT individuals were found in the bilateral superior and middle temporal gyri; middle and inferior frontal gyri; precentral gyrus; precuneus, posterior cingulate gyrus; intraparietal and occipital gyri; and the left superior frontal and postcentral gyri. VPT individuals also demonstrated widespread surface area reductions across the bilateral frontal, parietal, temporal, occipital, insular, and cingulate regions. In terms of cortical thickness, VPT individuals exhibited thinner cortices in the bilateral superior frontal, precentral, paracentral, postcentral, supramarginal, superior parietal, precuneus, and posterior cingulate regions, as well as the left banks of the superior temporal sulcus, middle frontal, and inferior parietal regions. Effect sizes of statistically significant cortical differences at each developmental stage are shown in Figure 3b.

**Figure 3.**
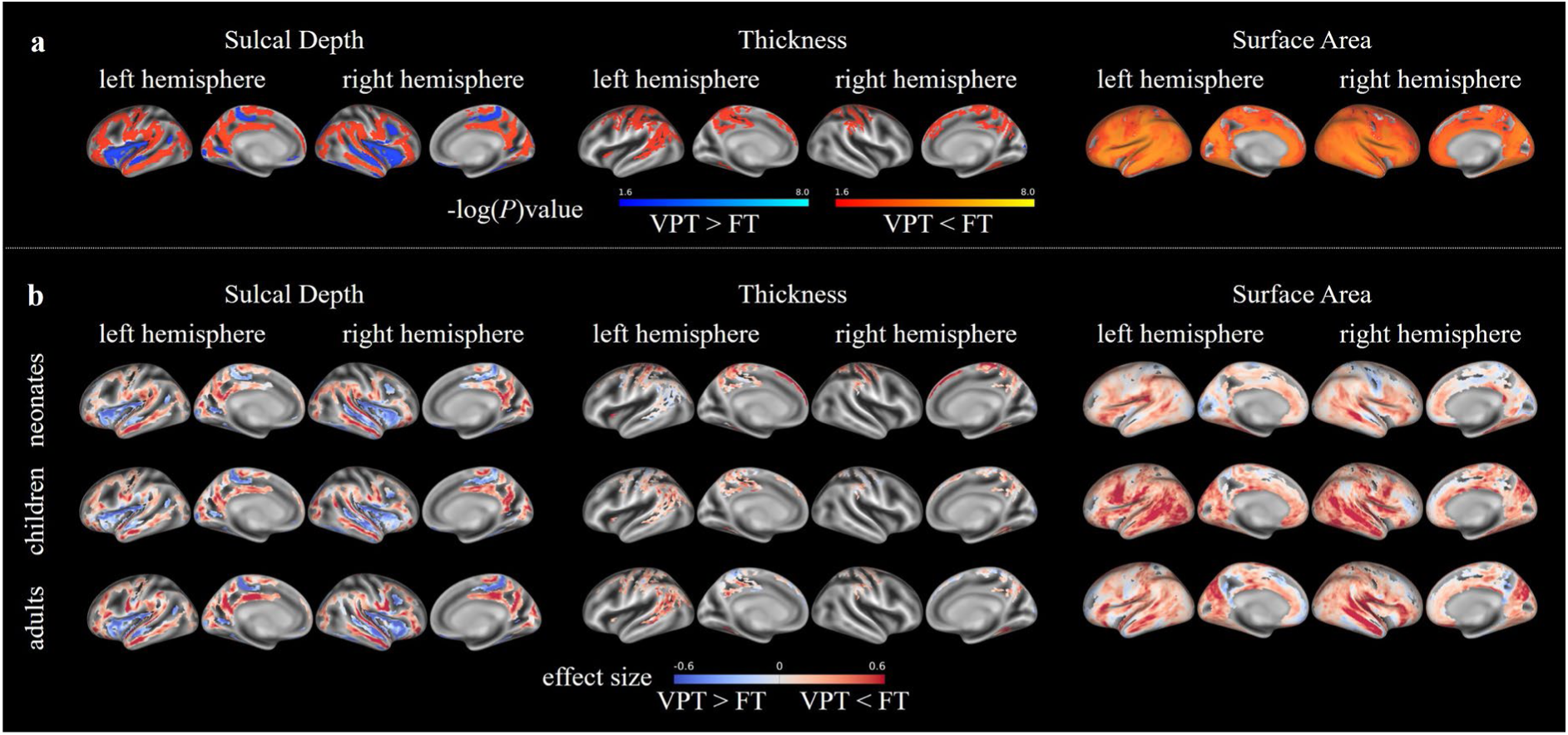
Group comparisons between very preterm (VPT) and full-term (FT) across development. **a)** Statistically significant cortical differences between VPT and FT groups across development. The blue–light blue colour scale represents higher values in VPT than in FT; the red-yellow colour scale represents reduced values in VPT compared with FT. **b)** Vertex-wise effect size within the significant regions identified in the cross-stage analyses for neonates, children and adults. Positive values (red) indicate VPT < FT, and negative values (blue) indicate VPT > FT.

Group comparisons of mean values within the significant regions identified in the cross-stage analyses are also presented in Table 2. Compared with full-term individuals, the VPT group exhibited smaller bilateral surface area and reduced cortical folding, consistently across development. In contrast, cortical thickness did not show a consistent pattern of group difference, with thinner bilateral cortex observed only in VPT neonates and thinner left-hemispheric cortex in VPT adults.

**Table 2.**
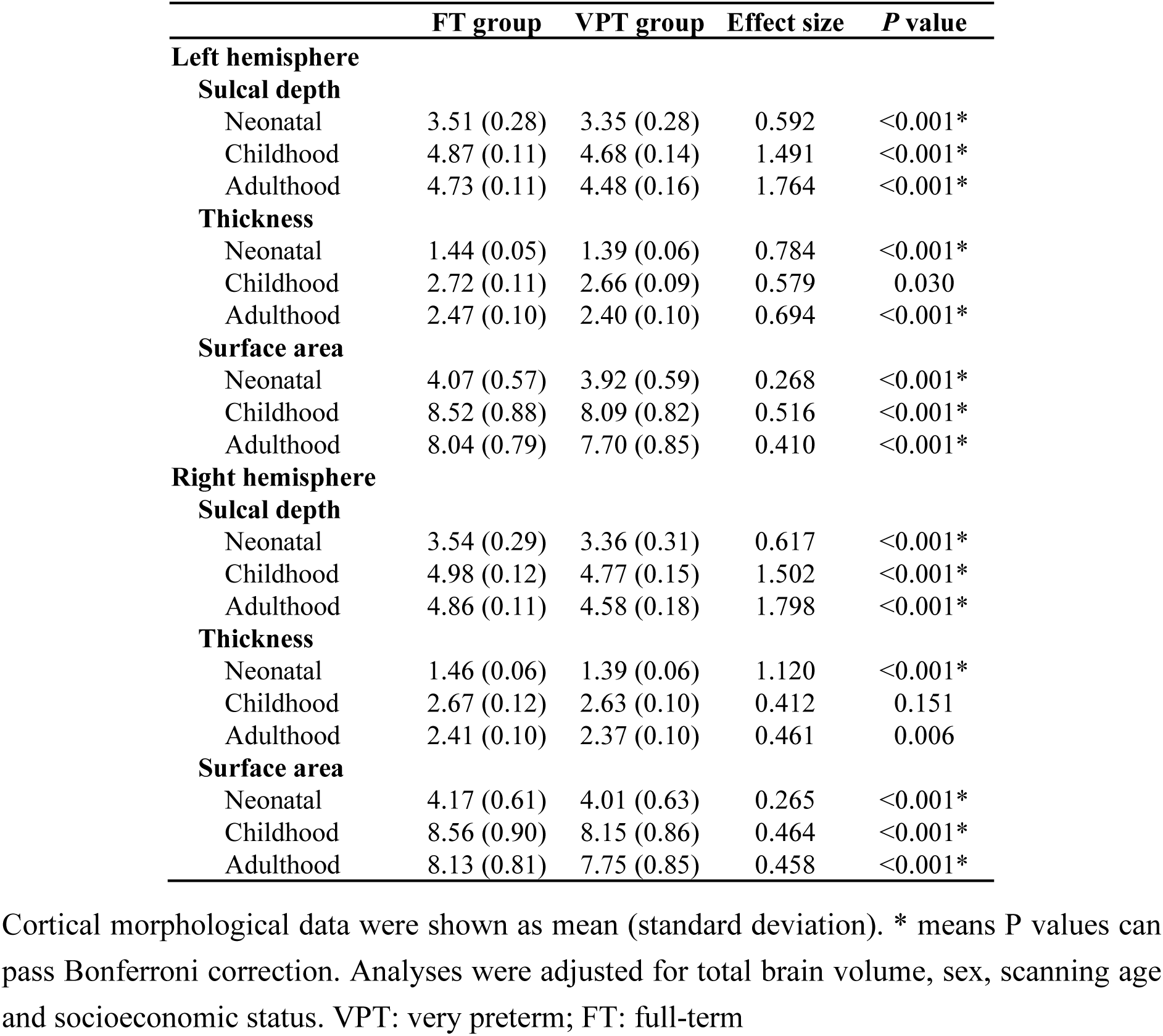
Group comparisons of mean values within the statistically significant regions between VPT and FT individuals.

|  | **FT group** | **VPT group** | **Effect size** | **P value** |
| --- | --- | --- | --- | --- |
| **Left hemisphere** |  |  |  |  |
| **Sulcal depth** |  |  |  |  |
| Neonatal | 3.51 (0.28) | 3.35 (0.28) | 0.592 | <0.001* |
| Childhood | 4.87 (0.11) | 4.68 (0.14) | 1.491 | <0.001* |
| Adulthood | 4.73 (0.11) | 4.48 (0.16) | 1.764 | <0.001* |
| **Thickness** |  |  |  |  |
| Neonatal | 1.44 (0.05) | 1.39 (0.06) | 0.784 | <0.001* |
| Childhood | 2.72 (0.11) | 2.66 (0.09) | 0.579 | 0.030 |
| Adulthood | 2.47 (0.10) | 2.40 (0.10) | 0.694 | <0.001* |
| **Surface area** |  |  |  |  |
| Neonatal | 4.07 (0.57) | 3.92 (0.59) | 0.268 | <0.001* |
| Childhood | 8.52 (0.88) | 8.09 (0.82) | 0.516 | <0.001* |
| Adulthood | 8.04 (0.79) | 7.70 (0.85) | 0.410 | <0.001* |
| **Right hemisphere** |  |  |  |  |
| **Sulcal depth** |  |  |  |  |
| Neonatal | 3.54 (0.29) | 3.36 (0.31) | 0.617 | <0.001* |
| Childhood | 4.98 (0.12) | 4.77 (0.15) | 1.502 | <0.001* |
| Adulthood | 4.86 (0.11) | 4.58 (0.18) | 1.798 | <0.001* |
| **Thickness** |  |  |  |  |
| Neonatal | 1.46 (0.06) | 1.39 (0.06) | 1.120 | <0.001* |
| Childhood | 2.67 (0.12) | 2.63 (0.10) | 0.412 | 0.151 |
| Adulthood | 2.41 (0.10) | 2.37 (0.10) | 0.461 | 0.006 |
| **Surface area** |  |  |  |  |
| Neonatal | 4.17 (0.61) | 4.01 (0.63) | 0.265 | <0.001* |
| Childhood | 8.56 (0.90) | 8.15 (0.86) | 0.464 | <0.001* |
| Adulthood | 8.13 (0.81) | 7.75 (0.85) | 0.458 | <0.001* |

Cortical morphological data were shown as mean (standard deviation). \* means P values can pass Bonferroni correction. Analyses were adjusted for total brain volume, sex, scanning age and socioeconomic status. VPT: very preterm; FT: full-term

### 3.3 Associations between clinical factors, neurodevelopmental outcomes and cortical alterations

Given the absence of consistent cortical thickness changes across developmental stages, only vertex-wise surface area and sulcal depth values from significant regions identified across development were included as cortical variables in the sCCA analyses. In the two infant cohorts, the sCCA analysis included 7,252 vertex-wise cortical features and two clinical variables (gestational age and birth weight). The corresponding sCCA results are shown in Figure 4. In dHCP, two canonical correlation modes survived Bonferroni correction and captured the same clinical-cortical associations: higher GA and BW were associated with greater cortical surface area and folding in frontal, temporal, parietal, and insular regions (*r*=0.753, *P_perm_*<0.001), whereas lower GA and BW were associated with reductions in both cortical surface area and folding (*r*=0.715, *P_perm_*<0.001). In ePrime, sCCA identified one canonical correlation mode which survived Bonferroni correction, linking greater GA and BW to higher brain scores, reflecting larger surface areas and increased cortical folding in frontal, temporal, parietal, and insular regions (*r*=0.745, *P_perm_*<0.001), consistent with the dHCP findings. Bootstrapping results implied the robustness of these findings, as the 95% confidence intervals of the resampling distribution did not include zero (Supplementary Figure 3).

**Figure 4.**
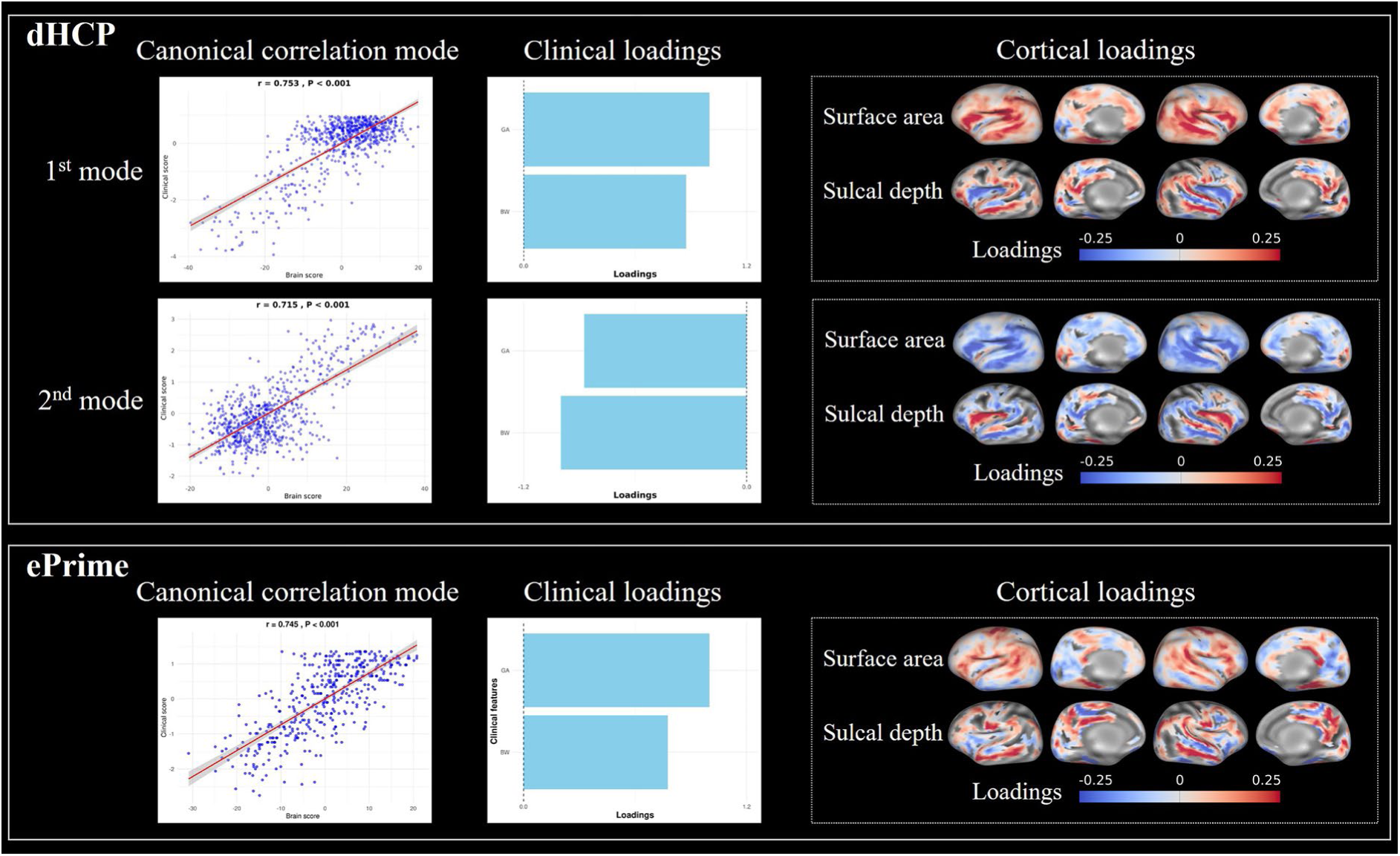
Associations between neonatal clinical factors and cortical variables identified by sCCA in dHCP and ePrime. Two significant canonical correlation modes were identified in dHCP (the first and second rows); and one significant mode was found in ePrime (the third row). Each significant canonical correlation mode comprised a pair of canonical variates derived from the CCA model: a clinical score, representing the canonical variate of the clinical variables, and a brain score, representing the canonical variate of the vertex-wise cortical variables. Clinical loadings were defined as the correlations between each clinical variable and the clinical score, whereas cortical loadings were defined as the correlations between each vertex-wise cortical variable and the brain score. Clinical factors included gestational age (GA) and birth weight (BW) in two infant cohorts. Cortical variables included 7,252 vertex-wise surface area and sulcal depth values derived from regions showing significant group differences across development.

Associations between brain scores and neurodevelopmental outcomes in two infant cohorts are shown in Figure 5. In dHCP, no significant group × brain scores interactions with neurodevelopmental outcomes were observed; therefore, analyses were performed across preterm and full-term participants. Brain scores from both modes showed significant associations with cognitive (*r*=0.133, *P*=0.002; *r*=-0.129, *P*=0.003) and motor outcomes (*r*=0.179, *P*<0.001; *r*=-0.172, *P*<0.001) after Bonferroni correction, indicating that greater cortical surface area and folding were associated with better cognitive and motor function. In ePrime, the brain score, driven by greater surface area and cortical folding, was positively correlated with cognitive scores (*r*=0.110, *P*=0.035), but this association did not survive Bonferroni correction. In contrast, the correlation with motor scores (*r*=0.191, *P*<0.001) remained significant after Bonferroni correction. No significant associations with language scores after Bonferroni correction were observed in either cohort.

**Figure 5.**
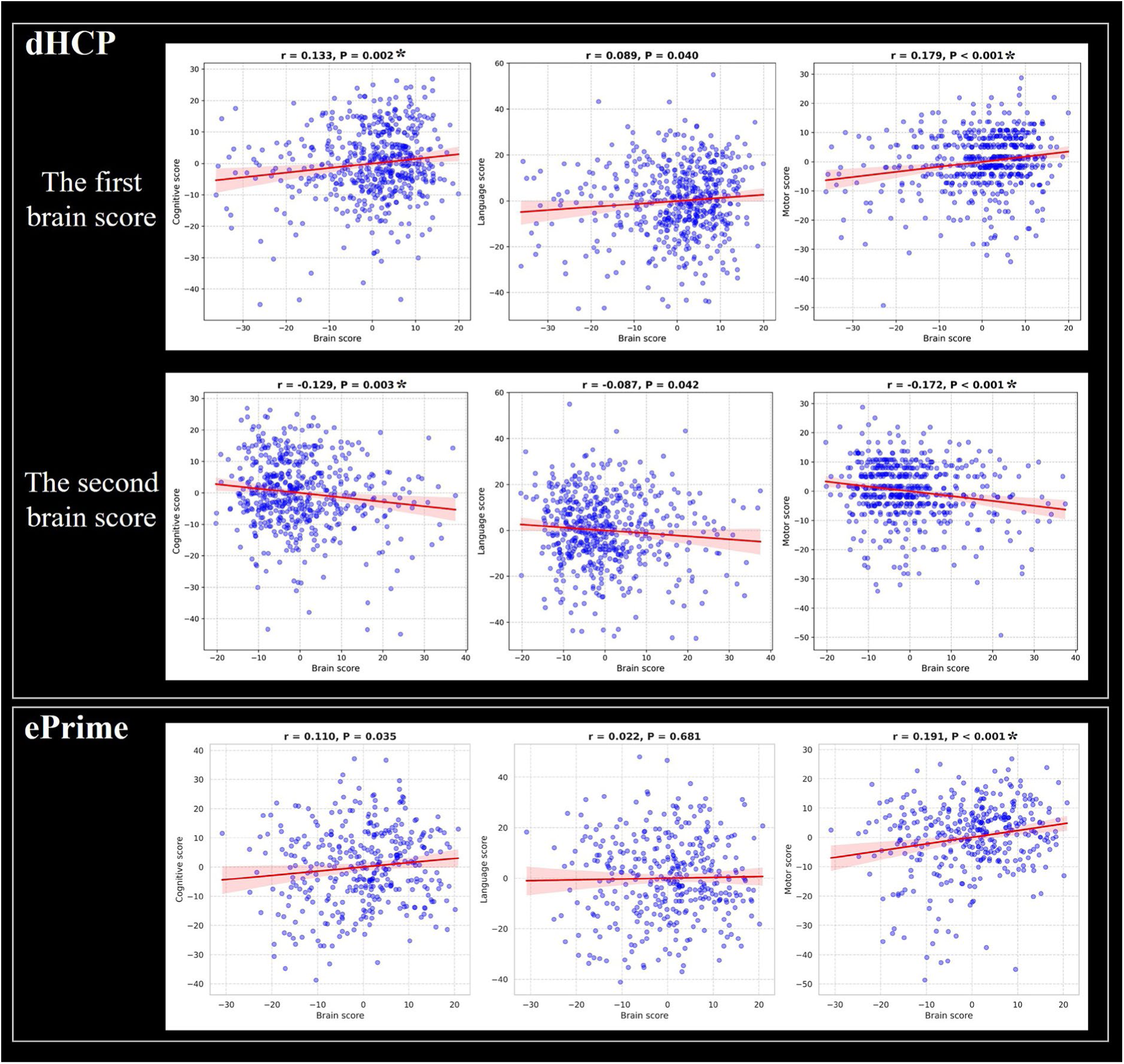
Associations between brain scores at term-equivalent age (37–45 weeks postmenstrual age) and neurodevelopmental outcomes at 18-24 months in dHCP and ePrime. In dHCP, correlations between brain scores from the two significant modes and cognitive, language, and motor scores are shown in the first and second rows, respectively. In ePrime, correlations between the brain score from the significant mode and cognitive, language, and motor scores are shown in the third row. * P < 0.05 after Bonferroni correction.

To externally validate the identified canonical patterns, the brain and clinical loadings from the two significant canonical variates derived from the dHCP sCCA model were projected onto the ePrime data to generate corresponding brain and clinical scores. The associations did not change substantially, with brain scores remaining significantly associated with motor outcomes (r=0.187, *P*<0.001; r=−0.185, *P*<0.001), whereas no significant associations were observed with cognitive or language outcomes. Reciprocal validation applying canonical loadings derived from the ePrime to the dHCP data yielded similar findings, with a significant association between brain scores and motor outcomes only (r=0.114, *P*=0.008) after Bonferroni correction (Supplementary Figure 4).

### 3.4 Associations of cortical expansion from infancy to childhood with GA and childhood IQ within VPT participants

Figure 6a shows the average relative cortical surface area expansion map across 148 VPT participants. The spatial pattern of cortical surface area expansion showed relatively low expansion in primary sensorimotor regions and high expansion in association cortices, consistent with the hierarchical sequence of brain maturation in which primary sensorimotor regions mature earlier than association cortices (Sydnor et al., 2021). Figures 6b and 6c show the associations of cortical expansion from infancy to childhood with GA and childhood IQ, respectively, in VPT participants. Greater bilateral cortical expansion was significantly associated with higher childhood full-scale IQ (left: r = 0.356, P < 0.001; right: r = 0.344, P < 0.001), although not with GA (left: r = −0.064, P = 0.441; right: r = −0.069, P = 0.406).

**Figure 6.**
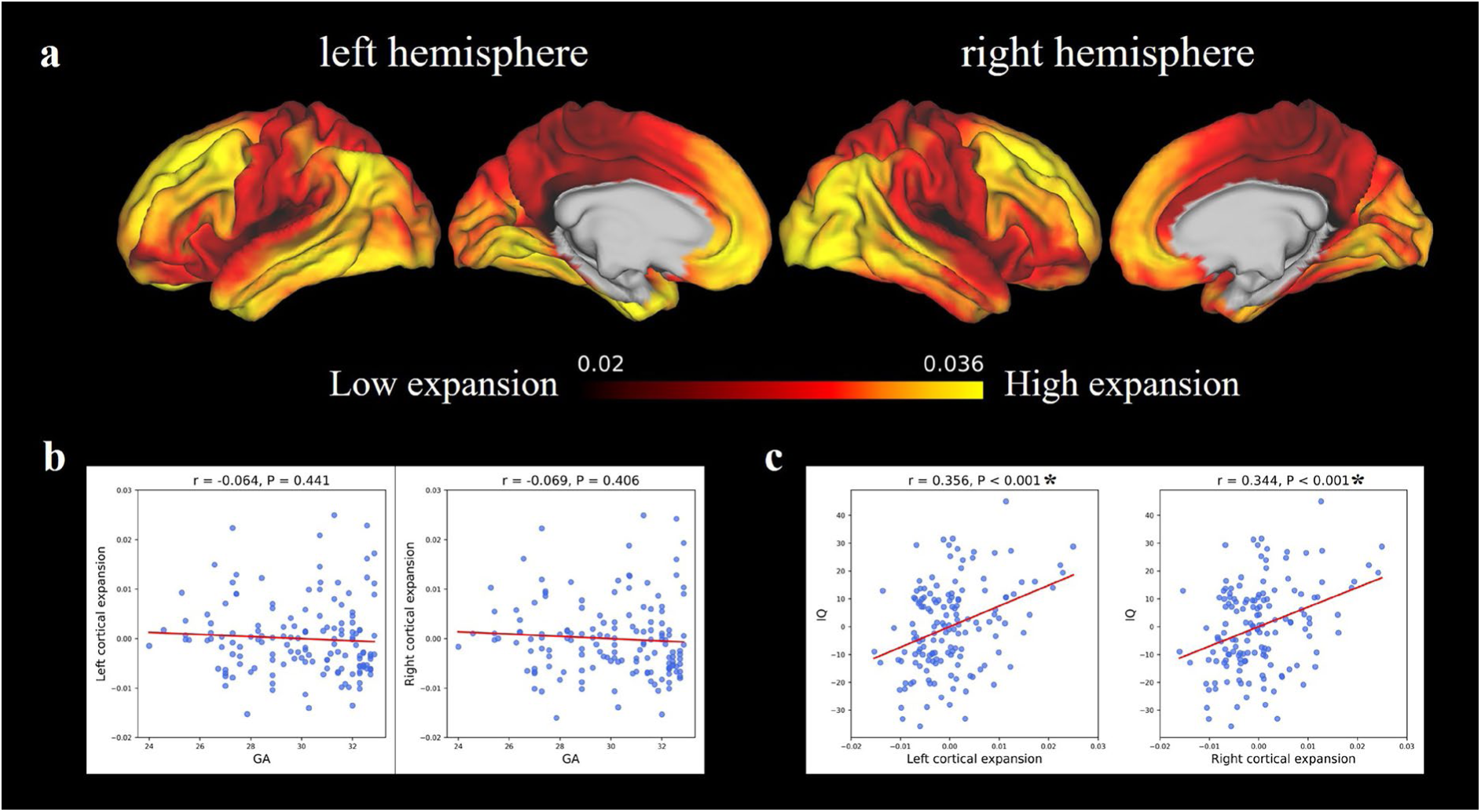
Longitudinal cortical expansion analyses in VPT participants. **(a)** Average cortical expansion map from the neonatal period to childhood. **(b)** Associations between mean cortical expansion within regions showing significant group differences and gestational age (GA). **(c)** Associations between mean cortical expansion within regions showing significant group differences and childhood IQ. * P < 0.05 after Bonferroni correction.

## 4 Discussion

In this study, we identified a cortical signature of VPT birth, characterized by smaller surface areas and reduced cortical folding in frontal, temporal, parietal and insular regions, consistently observed across cross-sectional cohorts spanning different developmental stages. We further found that the cortical footprint in neonates was associated with clinical factors and poorer motor function at 18-24 months and that greater cortical expansion from the neonatal period to childhood was associated with higher childhood full-scale IQ. Together, these findings highlight the persistence of cortical morphological alterations following very preterm birth and suggest that both neonatal cortical features and subsequent patterns of cortical expansion are associated with neurodevelopmental outcomes.

Overall, our findings were more stable across development for surface area and sulcal depth. By contrast, cortical thickness alterations varied across developmental stages, displaying both regional increases and decreases in neonates, no significant between-group differences in childhood, and cortical thinning of the lateral temporal regions in adults. It is possible that these findings point to distinct neurobiological mechanisms. Previous research has suggested that early-life stressors, including VPT birth, may be linked to accelerated cortical thinning and brain aging, further leading to mild cognitive impairments in later adulthood (Karolis et al., 2017; Kelly et al., 2023; Tooley et al., 2021). Notably, our findings in neonates and children contrast with previous findings, suggesting all results should be reviewed with caution. Previous research has indicated that VPT neonates display globally increased cortical thickness (Dimitrova et al., 2021; Thalhammer et al., 2024). During childhood, studies have reported heterogeneous patterns of cortical thickness alterations, including thinning in the lateral frontal and temporal lobes, increased thickness in occipital and frontal areas alongside thinning in lateral parietal and posterior temporal regions, and limited associations between cortical thickness and GA (El Marroun et al., 2020; Sripada et al., 2018; Thalhammer et al., 2024). These findings are likely due to the potential individual heterogeneity in cortical thickness (Thalhammer et al., 2024), differences in image processing pipelines, and the sensitivity of cortical thickness measures to image quality, particularly in neonatal MRI. Smaller brain size, lower spatial resolution due to shorter scan times, and increased motion artifacts exacerbate partial volume effects in neonatal MRI, reducing the accuracy of cortical thickness measurements (Makropoulos et al., 2018). Therefore, given the lack of consistent cortical thickness findings across development and its sensitivity to methodological variability, cortical thickness was excluded from our association analyses.

Our findings of reduced surface area in the frontal, temporal, and parietal regions in VPT individuals across development are consistent with previous studies (Engelhardt et al., 2015; Kelly et al., 2023; Kline et al., 2019; Thalhammer et al., 2024; Zhang et al., 2015). Furthermore, our findings indicated that VPT individuals exhibit shallower sulci and flatter gyri in these regions, suggesting that the observed widespread reductions in surface area might be driven, at least in part, by reduced cortical folding. We also identified cortical alterations across development in the insula, a finding that has been reported less frequently in previous studies (Inder et al., 2023; Kelly et al., 2023). This discrepancy may reflect differences in the GA distribution across study cohorts. Unlike the more complex secondary and tertiary cortical folding, the insular cortex begins to develop at approximately 15 weeks of gestation and undergoes earlier maturation compared to other regions (Kostovic et al., 2019). Consequently, insular development may be relatively preserved in cohorts including a greater proportion of later-born preterm infants, whereas alterations may be more evident in cohorts enriched for VPT births.

In both infant cohorts, we observed associations between cortical alterations linked to neonatal clinical factors and motor scores in toddlerhood, after accounting for sex and socioeconomic status. These findings support the importance of greater surface areas and cortical folding in the frontal, parietal, temporal, and insular regions for better neurodevelopmental outcomes, potentially reflecting enhanced global network efficiency (Kim et al., 2014). Importantly, the cortical markers in our study were characterized by extensive alterations in surface area and cortical folding, supporting the view that behavioural functions rely on integrated brain networks rather than isolated regions (Kim et al., 2014). Early motor development is known to play a critical role in shaping infants’ interactions with their environment, laying the foundation for subsequent cognitive and social-emotional development (Houwen et al., 2016). Impairments in these domains are commonly observed in neurodevelopmental conditions, such as intellectual disabilities, autism and ADHD (Purugganan, 2018; Shaw, 2024). Additionally, in childhood and beyond, the frontal, parietal, temporal, and insular cortices are central to cognitive, emotional, and behavioural regulation (Inder et al., 2023), and their morphological abnormalities have been implicated in a range of neuropsychiatric disorders, particularly those implicating affective and cognitive functions (Matsumoto et al., 2023). These findings may provide insights into the neurobiological mechanisms underlying the increased risk of neurodevelopmental difficulties in individuals born VPT.

We did not observe significant associations between cortical markers and language scores after adjusting for sex and socioeconomic status in either cohort. These findings are consistent with previous research, suggesting that language development may be more strongly shaped by a child’s postnatal environment context rather than by neonatal risk factors associated with VPT birth (Lean et al., 2018; Panceri et al., 2020). Because language acquisition critically depends on the quality and quantity of linguistic exposure and caregiver-child interactions, it may be particularly susceptible to environmental influences (De Leeuw et al., 2024; Houwen et al., 2016).

Notably, within the longitudinal VPT cohort, greater cortical expansion from the neonatal period to childhood in frontal, parietal, temporal and insular regions was associated with higher childhood full-scale IQ, highlighting the importance of postnatal cortical maturation for later cognitive development. In contrast, cortical expansion was not significantly associated with GA at birth. One possible explanation is that, once birth occurs during the third trimester of gestation, cortical maturation follows a broadly similar developmental trajectory, with the initial disruption associated with VPT delivery exerting a greater influence on cortical maturation than differences in GA. Alternatively, GA may be an imprecise marker of the biological processes underlying cortical development, acting instead as a proxy for a range of prenatal, perinatal and postnatal factors, including maternal characteristics, neonatal complications, and the family and school environment, all of which may differentially influence cortical maturation (Inder et al., 2023; Luby et al., 2023; Nawa et al., 2020; Silver et al., 2022). Together, these findings suggest that variability in cortical maturation within VPT individuals is likely driven by a combination of biological and environmental factors that are not fully captured by GA alone. Identifying the factors that promote cortical expansion may therefore provide opportunities for targeted interventions to improve cognitive outcomes in individuals born very preterm.

Although this study provides a systematic investigation of cortical alterations in VPT individuals, several limitations should be noted. First, group comparisons across different developmental stages were based on cross-sectional datasets. Although longitudinal data were available for one VPT cohort, corresponding longitudinal data for full-term individuals were unavailable. Therefore, the present study could not directly compare cortical developmental trajectories between VPT and full-term individuals, and altered postnatal cortical developmental trajectories following VPT birth cannot be excluded. Future longitudinal studies including both VPT and full-term cohorts are needed to validate these findings. Additionally, while GA did not differ significantly between VPT participants across datasets, differences in other unmeasured clinical characteristics may have contributed to variability between datasets. The lack of harmonized clinical information (e.g. singleton status and neonatal complications) limited our ability to account for these potential confounders in the analyses. Future research should investigate how perinatal factors influence cortical development in VPT individuals to inform early intervention strategies. Secondly, although widespread shallower sulci and flatter gyri were observed in VPT participants across development, separate vertex-wise analyses of each developmental stage yielded fewer significant findings. This may be due to limited statistical power or the limitations of diffeomorphic image registration, which cannot completely account for heterogeneity of cortical shape across individuals (Guo et al., 2025). When applying false discovery rate (FDR) correction (Noble et al., 2022), a less conservative approach than FWE correction, we observed widespread shallower sulci and flatter gyri at each developmental stage (Supplementary Figure 5).

Finally, associations between cortical markers and neurodevelopmental outcomes showed small effect sizes. One recent study suggested that low reliability across behavioural phenotypes can attenuate or obscure true brain-behaviour associations (Gell et al., 2024). While our study identified correlations between neonatal clinical factors, neurodevelopmental outcomes and cortical alterations, the complex interplay and potential causal relationships among these factors remain challenging to elucidate. It is widely recognized that neuroimaging data alone may have limited predictive power for behavioural outcomes (Zhi et al., 2024). Integrating additional data types, including multiple environmental exposures and more detailed clinical measures, may improve predictive accuracy in future studies.

In conclusion, by integrating data across neonatal, childhood, and adulthood stages, we identified a consistent VPT cortical signature characterized by reduced surface area and cortical folding in the frontal, temporal, parietal, and insular regions. This signature was associated with neonatal clinical factors and poorer motor outcomes at 18–24 months. Within VPT individuals, greater cortical expansion of this signature from the neonatal period to childhood was associated with higher childhood full-scale IQ, indicating the importance to study both neonatal cortical features and subsequent cortical expansion in relation to neurodevelopmental outcomes. Future research should further investigate the genetic and environmental mechanisms underlying the cortical signature to advance our understanding of the origins of neurodevelopmental disorders and inform more targeted interventions.

## Declaration of the use of AI

AI (or other tools) were used to check spelling and grammar.

## Data and Code availability

The dHCP data are available at https://biomedia.github.io/dHCP-release-notes/. The ePrime, BIPP and UCLH cohort datasets that support the findings of this study are available from the corresponding author upon reasonable request. The dHCP deep learning-based pipeline: https://github.com/m-qiang/dhcp-dl-neonatal.git. The dHCP spatiotemporal cortical surface templates that are compatible with cortical surface outputs from deep learning-based pipeline: https://github.com/kaili23/VPT_corticalAlterations/tree/main/dHCP/templates. MSMSulc configuration: https://github.com/kaili23/VPT_corticalAlterations/blob/main/Configs. Code used in this manuscript is available at https://github.com/kaili23/VPT_corticalAlterations.

## Acknowledgements

We would like to thank all the participants and their families who were involved in the dHCP, ePrime, BIPP and UCLH cohorts. We are also grateful to all the researchers involved in the design, implementation, and recruitment for these four studies. The dHCP project was funded by the European Research Council (ERC) under the European Union Seventh Framework Programme (FR/2007-2013)/ERC grant agreement no. 319,456. The ePrime data were acquired during independent research funded by the National Institute for Health Research (NIHR) under its Programme Grants for Applied Research Programme (Grant Reference No. RP-PG-0707-10154). The BIPP study was supported by the Medical Research Council (UK) (grant no’s MR/S026460/1 [to C.N., A.D.E., S.J.C.]). The UCLH study was funded by the Medical Research Council, UK (Grant No. MR/K004867/1 [to C.N.]). We acknowledge use of the King’s Computational Research, Engineering and Technology Environment (CREATE) (https://doi.org/10.18742/rnvf-m076) and infrastructure support from the NIHR Mental Health BRC at South London and Maudsley NHS Foundation Trust, King’s College London.

K.L. is supported by NIHR Maudsley Biomedical Research Centre (BRC) PhD studentship.

E.C.R has been supported by funding from the MRC methodology grant (MR/V03832X/1 and UKRI534) and the Medical Research Council Centre for Neurodevelopmental Disorders,

King’s College London (MR/N026063/1). The views expressed are those of the authors and not necessarily those of the NHS, the NIHR or MRC.

## Author contribution

K.L., E.C.R., and C.N. designed the research. K.L., Y.G., L.Z.J.W., R.B., Z.S., M.L. and M.Q. performed research and analysed data. K.L., D.R., P.D., A.D.E., E.C.R., and C.N. wrote and revised the manuscript. All authors reviewed and edited the manuscript.

## Supplementary material

Supplementary materials associated with this article are attached.

