## Supplementary for "A cortical signature of very preterm birth across development and its association with neurodevelopmental outcomes"

**Supplementary material**

**Participants**

**dHCP:** The dHCP recruited infants born between 23 and 44 week’s gestation at St Thomas’ Hospital, London, and imaged at the Evelina Newborn Imaging Centre, King's College London, between 2015 and 2019 (Edwards et al., 2022). Ethical approval was granted by the London Riverside Research Ethics Committee of the Health Research Agency (REC: 14/Lo/1169), and written informed consent was obtained from participants’ guardians. In this study, we included full-term neonates born at 37–42 weeks’ gestation and very preterm neonates born before 33 weeks’ gestation. All infants underwent MRI at term-equivalent age (37-45 weeks’ postmenstrual age). Exclusion criteria were contraindications to MRI, major congenital abnormalities (e.g., congenital cardiac anomalies), missing or corrupted structural MRI data, and failed image reconstruction.

**ePrime:** The ePrime study recruited VPT infants born before 33 weeks of gestation from 14 hospitals within the North and South-West London Perinatal Network between April 2010 and July 2013 (Edwards et al., 2018). The study was approved by the Hammersmith and Queen Charlotte’s Research Ethics Committee (REC: 09/H0707/98), and written parental consent was obtained for each infant before imaging. Inclusion criteria included birth before 33 weeks of gestation, maternal age over 16 years, and mothers who were not hospital inpatients. Major congenital malformations, contraindications to MRI, non-English-speaking parents, or being subject to child protection proceedings were excluded.

**BIPP:** The BIPP study is a childhood follow-up of the ePrime study, comprising VPT participants and full-term controls recruited at St Thomas’ Hospital, London, between 2020 and 2025 (Sun et al., 2023). The study was approved by the Southeast Research Ethics Committee (REC: 19/LO/1940) and the Stanmore Research Ethics Committee (REC: 18/LO/0048), and written informed consent was obtained from participants’ guardians. Inclusion criteria for the VPT group were birth before 33 weeks’ gestation, whereas controls were born at full term (37–42 weeks’ gestation). Exclusion criteria included a history of neurological conditions (e.g., meningitis or cerebral infections), major congenital malformations, and contraindications to MRI.

**UCLH cohort:** This study followed VPT neonates born at the UCLH Neonatal Unit between 1979 and 1985 into middle adulthood (mean age: 30 years) (Papini et al., 2020). The study was approved by the South London and Maudsley Research Ethics Committee and the Psychiatry, Nursing and Midwifery Research Ethics Subcommittee at King’s College London (REC: PNM/12/13-10), and written consent was acquired for each participant. Inclusion criteria for the VPT group were birth before 33 weeks’ gestation, whereas controls were born at full term (37–42 weeks’ gestation). For all participants, exclusion criteria included any history of neurological complications, including meningitis, head injury, and cerebral infections, and contraindications to MRI.

**Clinical Features**

In dHCP, 520/541 (96.1%) full-term controls were singletons, with a mean (SD) birth weight of 3.38 kg (0.51) for the full-term group, while 58/81 (71.6%) VPT participants were singletons, with a mean (SD) birth weight of 1.20 kg (0.43) for the VPT group.

In BIPP, 108/158 (68.35%) VPT participants were singletons, with a mean (SD) birth weight of 1.32 kg (0.39) for the VPT group; singleton status and birth weight data were unavailable for full-term controls. Eight VPT participants (5.1%) had major perinatal brain lesions, assessed by one experienced neuroradiologist using MRI and defined as cystic periventricular leukomalacia, more than ten punctate white matter lesions, and/or grade III–IV germinal matrix haemorrhage.

In UCLH cohort, 91/118 (77.12%) VPT participants were singletons, with a mean (SD) birth weight of 1.32 kg (0.36); singleton status and birth weight data were unavailable for full-term controls. Twenty-six VPT participants (22.0%) had major brain injuries, assessed using cranial ultrasound and defined as grade III–IV periventricular haemorrhage with ventricular dilatation, as detailed in Nosarti et al. (Nosarti et al., 2012).

Available clinical information for the ePrime is summarized in the table below. Of the participants, 305 (73.7%) were singletons and 256 (61.8%) were delivered by caesarean section. Major brain lesions on MRI were defined as cystic periventricular leukomalacia, more than ten punctate white matter lesions, and/or grade III–IV germinal matrix haemorrhage (Ball et al., 2017).

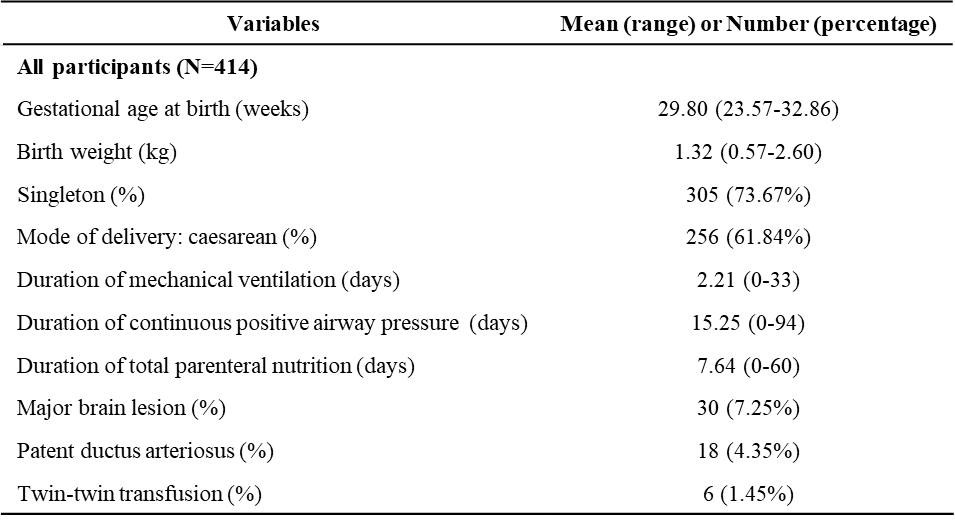

**MRI acquisition**

**dHCP:** MRI data for each neonate were acquired using a 3T Philips Achieva system equipped with a 32-channel coil at the Evelina Newborn Imaging Centre, Centre for the Developing Brain, King’s College London. Detailed MRI acquisition protocols have been previously described (Edwards et al., 2022). T2w images were obtained with a repetition time/echo time (TR/TE) of 12000/156 ms, captured in sagittal and axial slices with an in-plane resolution of 0.8mm×0.8mm and a slice thickness of 1.6 mm, overlapping by 0.8 mm. Motion correction and super-resolution reconstruction techniques were applied to produce isotropic volumes with a resolution of 0.5 mm.

**ePrime:** MRI data were acquired using a 3T Philips system with an 8-channel phased array head coil, located within the neonatal intensive care unit at Queen Charlotte’s and Chelsea Hospital, London. Scans were performed when infants were asleep, under the supervision of a skilled paediatrician. Infants whose parents chose sedation for the procedure (87%) received oral chloral hydrate (25–50 mg/kg) (Ball et al., 2017). T2-weighted images were obtained with the following parameters: TR/TE = 8670/160 ms, flip angle = 90°, field of view (FOV) = 220 x 220 mm, and matrix size = 256 x 256. The in-plane resolution was 0.86 × 0.86 mm, with a slice thickness of 2 mm and an overlap of 1 mm.

**BIPP:** MRI data were acquired using a Philips 3T Achieva system with a 32-channel head coil at the Evelina Newborn Imaging Centre, Evelina London Children’s Hospital. The study employed high-efficiency MR imaging techniques, including optimized rapid motion-tolerant acquisition and reconstruction methods, which significantly reduced scanning time and were therefore well-suited for pediatric participants. T1-weighted images were captured using a 3D MPRAGE sequence with the following parameters: TR/TE = 7.9/3.6 ms, flip angle = 8°, FOV = 240 × 220 × 160 mm, and voxel size = 1 mm isotropic.

**UCLH cohort:** MRI data were acquired using a 3T GE Signa HDx system with an 8-channel head coil at the Maudsley Hospital, London. T1-weighted images were acquired utilizing a fast spoiled gradient-echo sequence (FSPGR) sequence with the following parameters: TR/TE = 7.1/2.8 ms, flip angle = 20°, matrix = 256 × 256, and voxel size = 1.1 mm isotropic.

**Neonatal Template Generation**

Neonatal surface templates were generated using methods described in Bozek et al., 2018 and Williams et al., 2023 (Bozek et al., 2018; Williams et al., 2023). Firstly, each neonatal surface was aligned to the HCP template space using rigid alignment based on sulcal depth maps. Then week-specific sulcal depth templates were generated using an adaptive kernel weighting method proposed by Serag et al., (Serag et al., 2012) which uses a Gaussian kernel to define temporal weights ($w_{i}$) based on the difference between the subject’s scanning age ($t_{i}$) and the template age ($t$):

$$w_{i} (weights) = \frac{1}{\sigma\sqrt{2\pi}}exp\left( -\frac{1}{2}\left( \frac{t_{i} - t}{\sigma} \right)^{2} \right)$$

Doing this addresses the non-uniform distribution of gestational ages across different weeks, which can lead to inconsistencies in the anatomical detail retained in the weekly templates (Serag et al., 2012). Sigma ($\sigma$) was adaptively adjusted: smaller sigma values were used for weeks with more subjects, while larger sigma values were used for weeks with fewer subjects. This process for determining sigma involved first deriving the median number of subjects per week using a fixed kernel width (sigma = 1), which was then used as the target number of subjects per week. Subsequently, sigma was iteratively adjusted (increased or decreased) to ensure the sample size for each week reached this target number. To avoid excessively large sigma values, we set a tolerance threshold allowing sample sizes to vary within ±20 subjects of the target per week. In this study, the median target number was 58. The final number of subjects across the weeks (from 28 to 45 weeks) was as follows: 39, 39, 38, 38, 38, 40, 42, 64, 65, 50, 74, 78, 66, 69, 62, 70, 59, and 52. Corresponding sigma values were: 4.44, 3.44, 2.58, 1.86, 1.30, 1.01, 1.00, 1.00, 1.00, 1.00, 0.99, 0.57, 0.42, 0.28, 0.28, 0.42, 0.42, and 1 weeks.

Sulcal depth only captures coarse-scale folding patterns of the brain. Thus, templates were subsequently refined to align the more detailed patterns of gyrification and sulci captured by curvature maps. This process is iteratively repeated until the refinement of the curvature maps converges, ensuring that the templates can include the detailed anatomy of the brain. To remove the drift that occurred during the iterative registration process, the group average of inverse spheres was calculated for each week. This average sphere was then projected back onto the individual subjects’ surface sphere to obtain corrected surface spheres. Following the de-drifting, all templates of surface meshes and metrics were generated using the adaptive kernel weighting method.

To achieve left-right vertex correspondence, similar to HCP templates, MSMSulc was used to align the left and right sulcal depth maps for each gestational week (Williams et al., 2023). Registration was performed in both directions, left-to-right and right-to-left, following the method described in previous studies (Garcia et al., 2018; Williams et al., 2023). The deformation maps from the left-to-right registration were averaged with the inverse of the right-to-left registration to produce an intermediate template space with left-right vertex correspondence. Surface meshes and cortical metrics for each week were then resampled onto this left-right vertex correspondence template surface. Finally, MSMSulc registration was applied across consecutive weekly templates (e.g., 28 to 29 weeks, 29 to 30 weeks) to generate a series of spherical deformations. These deformations were concatenated to enable direct mapping from any week-specific template to the week-40 template, thus providing a common template space for group comparisons.

**Surface Registration**

For neonatal surface registration, the native space of each subject was firstly aligned to their respective week-specific dHCP templates using MSMSulc. To align each subject to the common week-40 template space, the deformation from the native space to the week-specific template was concatenated with the deformation from the week-specific template to the week-40 template. This step ensures that all neonates are consistently aligned to the week-40 template for subsequent analyses. For children and adults, the native cortical surfaces were aligned to the HCP template space using MSMSulc registration.

To enable integrated cross-stage analysis across development, neonatal templates need to be aligned to the HCP adult templates. This was achieved by registering the dHCP week-40 template to the HCP template using MSMSulc, producing a template-to-template registration deformation, as done in Williams et al,.(Williams et al., 2023) The registration deformation from dHCP to HCP was concatenated with the deformation from native to week-40 dHCP template, resulting in a registration deformation from the native space to the HCP template for each subject.

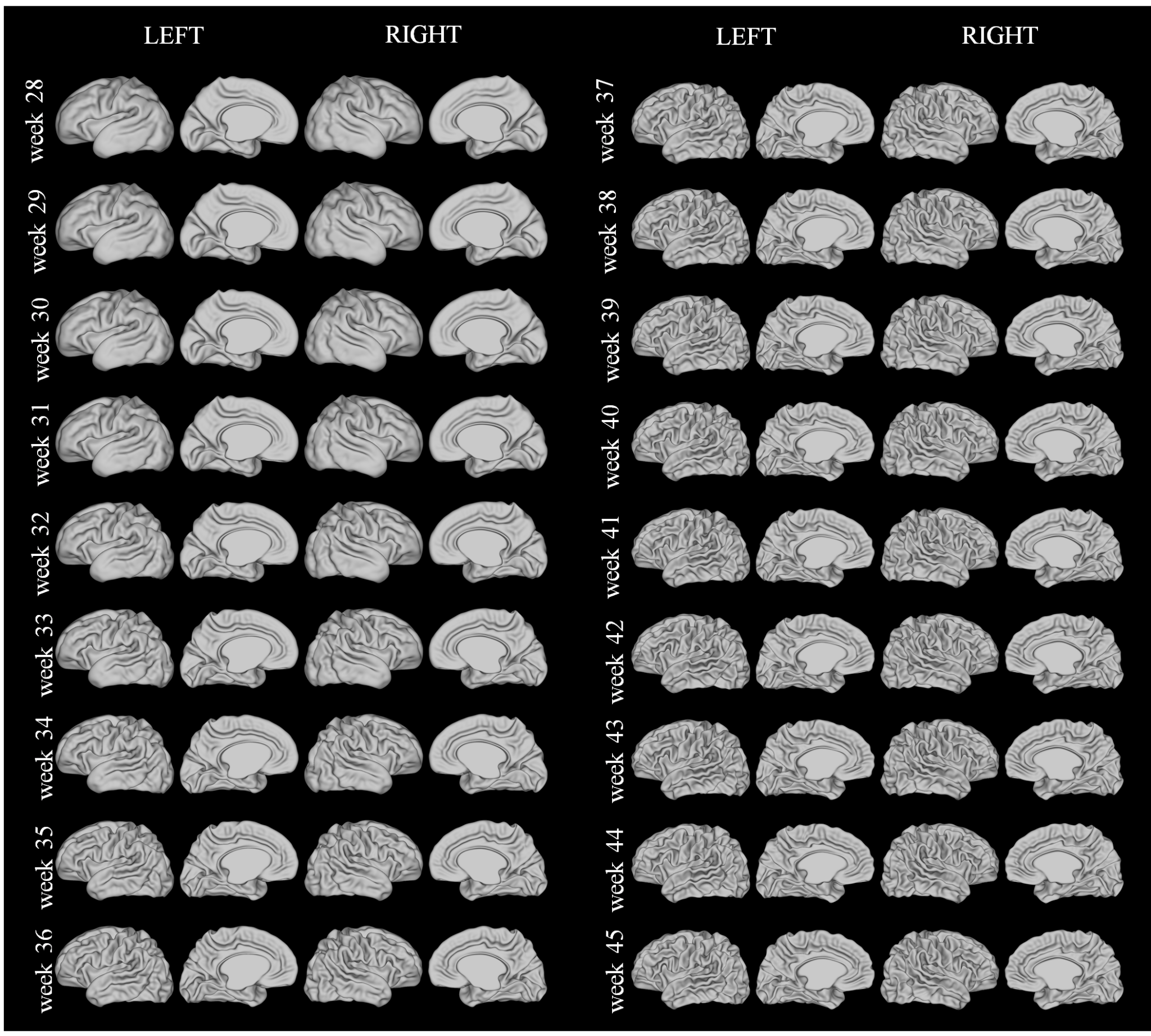

**Supplementary Figure 1. dHCP white matter surface templates for each week.**

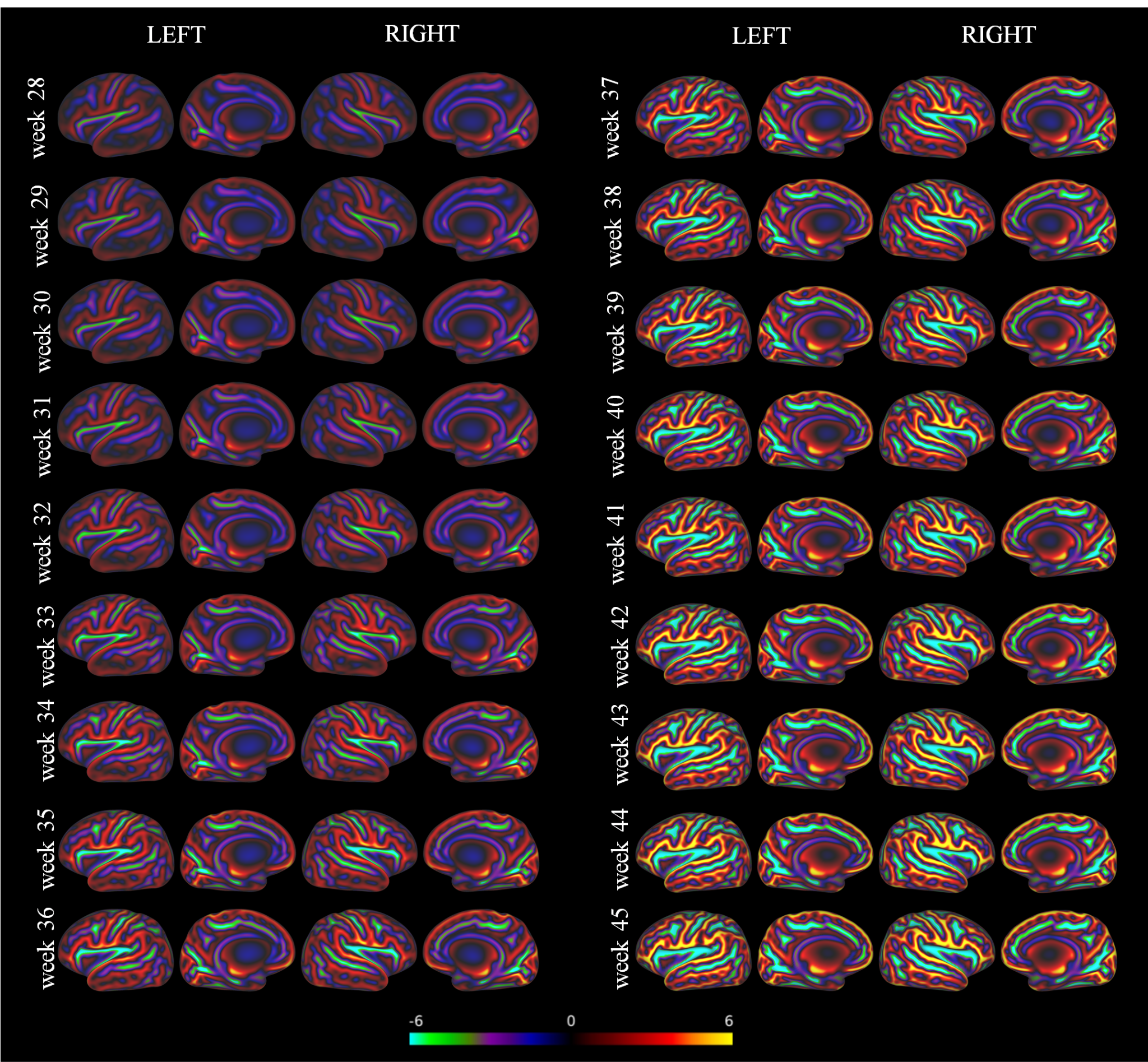

**Supplementary Figure 2. dHCP sulcal depth templates for each week.**

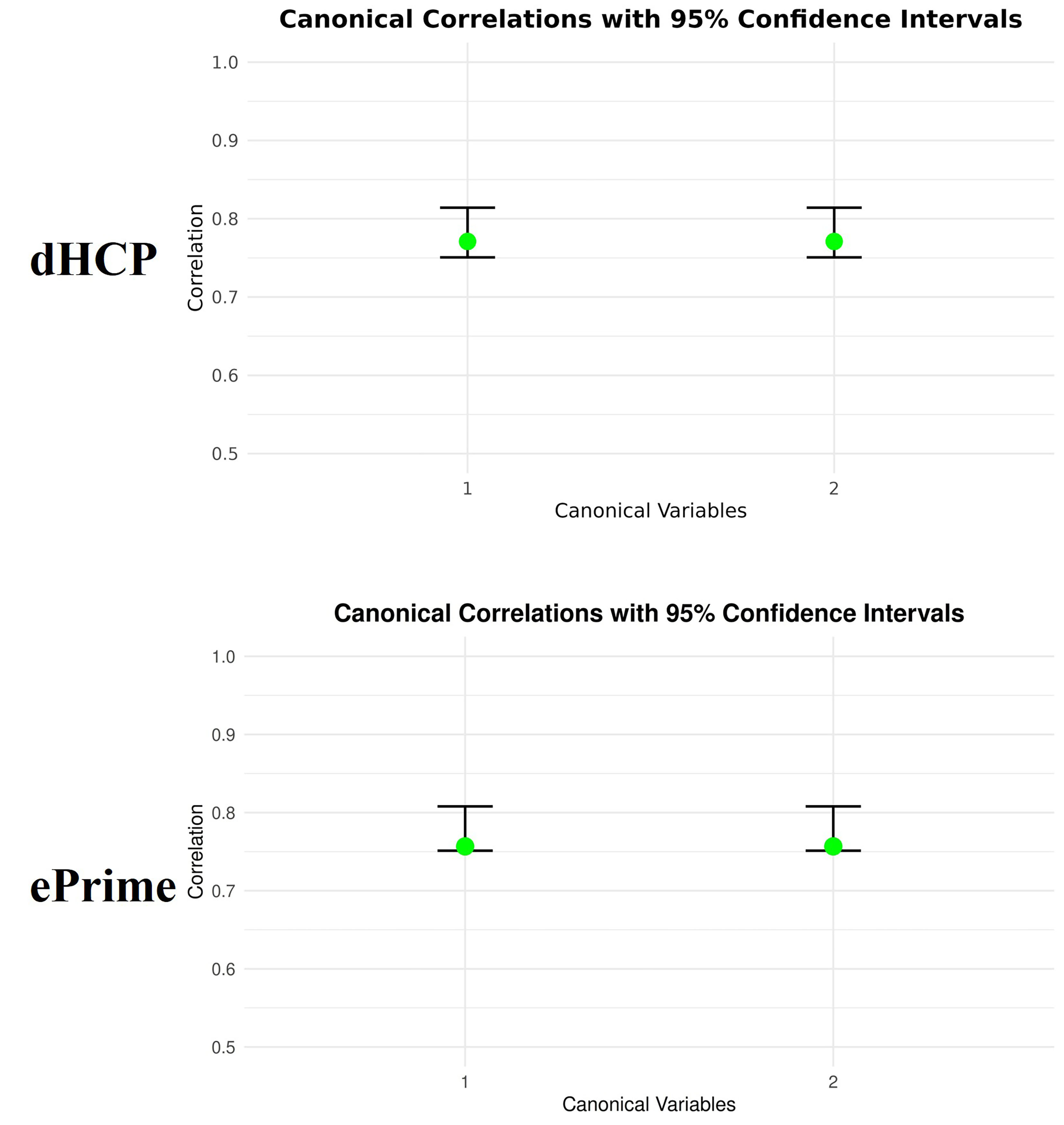

**Supplementary Figure 3. Bootstrapping results in dHCP and ePrime.** Resampling distribution of canonical correlations. Each bar represents the 95% confidence interval and each dot represents correlation coefficients in the original sample. The green dots demonstrate their 95% confidence intervals do not include zero.

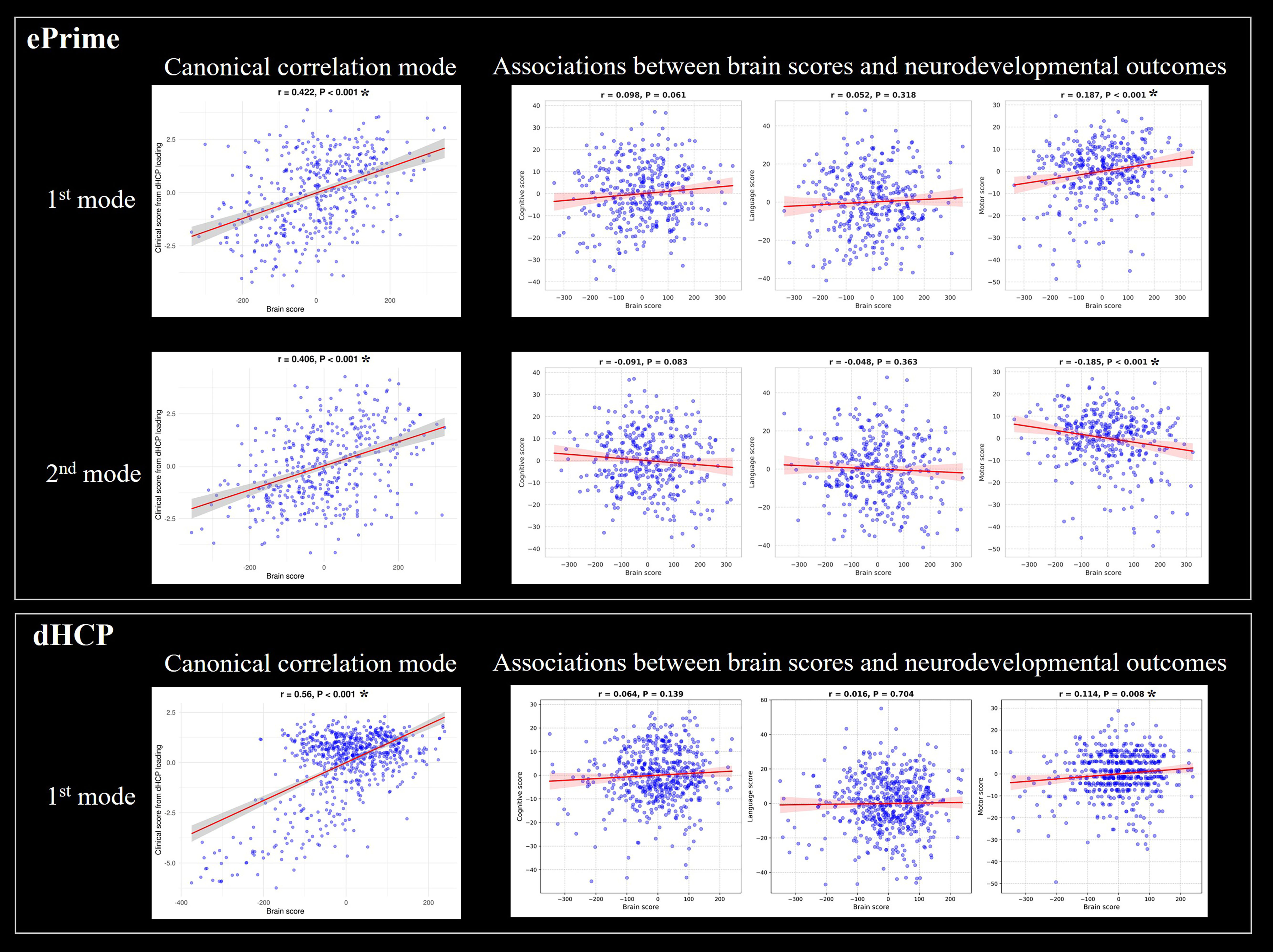

**Supplementary Figure 4. Cross-cohort validation of sparse CCA results in the two infant cohorts.** The first and second rows show the associations between brain scores and clinical scores, as well as between brain scores and neurodevelopmental outcomes, after applying the brain and clinical loadings from the two significant canonical variates derived from the dHCP sCCA model to the ePrime dataset. The third row shows the corresponding analyses after applying the canonical loadings derived from the ePrime to the dHCP cohort. * P < 0.05 after Bonferroni correction

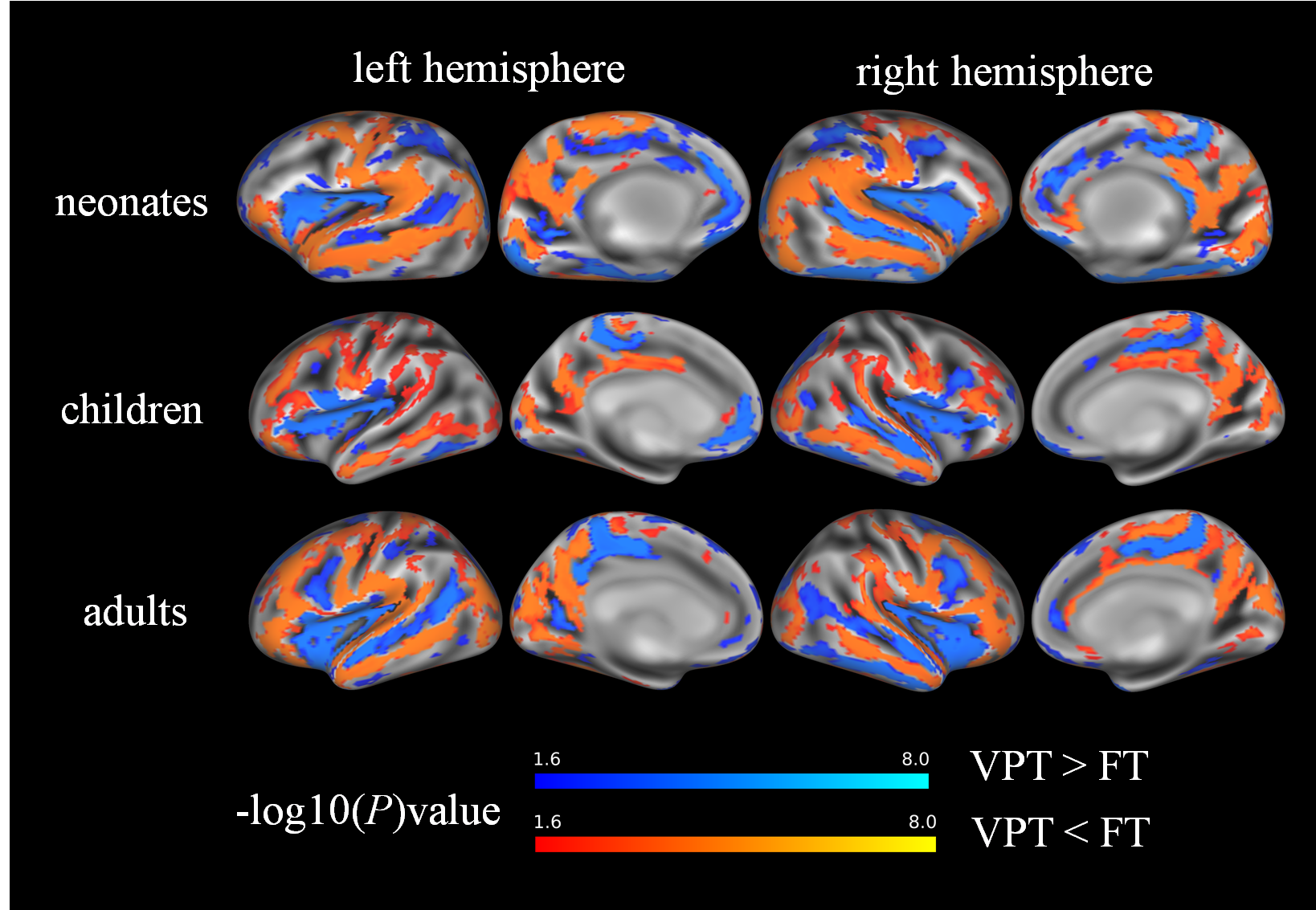

**Supplementary Figure 5.** Group differences in sulcal depth between very preterm (VPT) and full-term (FT) participants at each developmental stage. FDR (false discovery rate) correction was applied for multiple comparisons.

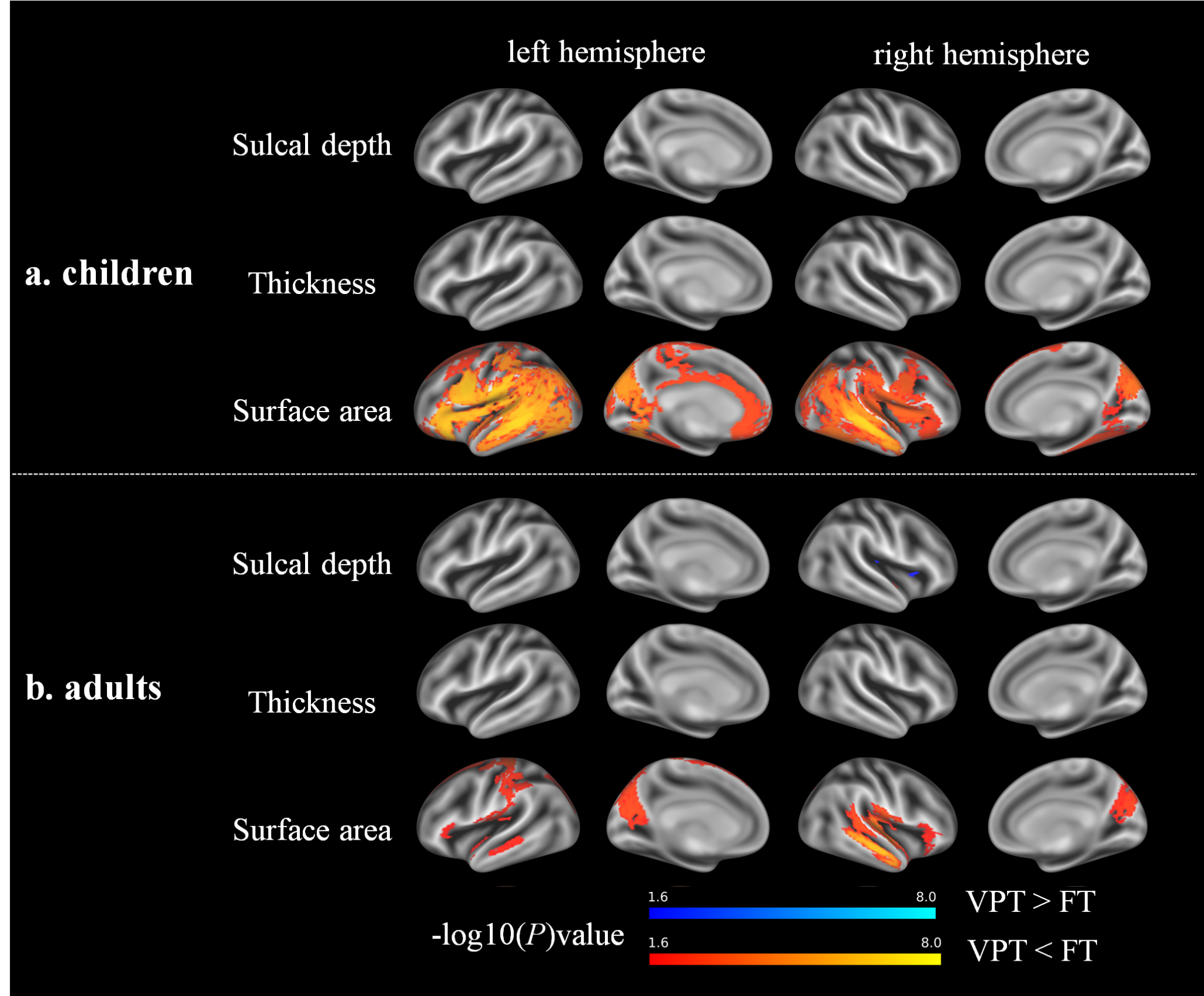

**Supplementary Figure 6.** Group differences between very preterm (VPT) and full-term (FT) participants after excluding VPT children (n=8; 5.06%) and adults (n=26; 22.03%) with major perinatal brain injuries. Widespread surface area reductions in the VPT group observed in the full sample remained largely unchanged.
